# Multi-domain touchscreen-based cognitive assessment of C57BL/6J female mice shows whole body exposure to ^56^Fe particle space radiation in maturity improves discrimination learning yet impairs stimulus-response habit learning

**DOI:** 10.1101/2021.06.09.447537

**Authors:** Ivan Soler, Sanghee Yun, Ryan P. Reynolds, Cody W. Whoolery, Fionya H. Tran, Priya L. Kumar, Yuying Rong, Matthew J. DeSalle, Adam D. Gibson, Ann M. Stowe, Fred C. Kiffer, Amelia J. Eisch

**Affiliations:** University of Pennsylvania Perelman School of Medicine, Philadelphia, PA USA; Department of Psychiatry, University of Texas Southwestern Medical Center, Dallas, TX, USA; Department of Anesthesiology and Critical Care Medicine, Children’s Hospital of Philadelphia, Philadelphia, PA USA; University of Pennsylvania, Philadelphia, PA USA; Department of Neurology and Neurological Therapeutics, University of Texas Southwestern Medical Center, Dallas, TX, USA; Department of Neuroscience and Mahoney Institute for Neurosciences, University of Pennsylvania Perelman School of Medicine, Philadelphia, PA USA

**Keywords:** dentate gyrus, prefrontal cortex, striatum, hippocampus, behavioral pattern separation, rodent touchscreen

## Abstract

Astronauts during interplanetary missions will be exposed to galactic cosmic radiation, including charged particles like ^56^Fe. Preclinical studies with mature, “astronaut-aged” rodents suggest space radiation diminishes performance in classical hippocampal- and prefrontal cortex-dependent tasks. However, a rodent cognitive touchscreen battery unexpectedly revealed ^56^Fe radiation improves the performance of C57BL/6J male mice in a hippocampal-dependent task (discrimination learning) without changing performance in a striatal-dependent task (rule-based learning). As other preclinical work suggests the female rodent brain may be relatively resistant to charged particle-induced injury, and as the proportion of female vs. male astronauts is increasing, further study on how charged particles influence the touchscreen cognitive performance of female mice is warranted. We hypothesized that, similar to mature male mice, mature female C57BL/6J mice exposed to whole-body ^56^Fe irradiation (3 × 6.7cGy ^56^Fe over 5 days, 600MeV/n) would improve performance vs.Sham conditions in touchscreen tasks relevant to hippocampal and prefrontal cortical function (e.g. location discrimination reversal [LDR] and extinction, respectively). In LDR, ^56^Fe female mice more accurately discriminated two discrete conditioned stimuli relative to Sham mice, suggesting improved hippocampal function. However, ^56^Fe and Sham female mice acquired a new simple stimulus-response behavior and extinguished this acquired behavior at similar rates, suggesting similar prefrontal cortical function. Based on prior work on multiple memory systems, we next tested whether improved hippocampal-dependent function (discrimination learning) came at the expense of striatal rule-based learning (visuomotor conditional learning). Interestingly, ^56^Fe female mice took more days to reach criteria in this striatal-dependent rule-based test relative to Sham mice. Together, our data support the idea of competition between memory systems, as an ^56^Fe-induced decrease in striatal-based learning is associated with enhanced hippocampal-based learning. These data emphasize the power of using a touchscreen-based battery to advance our understanding of the effects of space radiation on mission critical cognitive function in females, and underscore the importance of preclinical space radiation risk studies measuring multiple cognitive processes, thereby preventing NASA’s risk assessments from being based on a single cognitive domain.

## 1. Introduction

As space agencies plan for impending interplanetary missions - such as to Mars - understanding potential hazards associated with galactic cosmic radiation (GCR) exposure becomes a priority (Vazquez, 1998; Schimmerling et al., 2003; Setlow, 2003; National Research Council et al., 2008; Chancellor et al., 2014; Cucinotta, 2015; Kokhan et al., 2016; Nelson et al., 2016; National Academies of Sciences, Engineering, and Medicine et al., 2018). GCR is composed of fast-moving low and high-(H) atomic number (Z) and high-energy (E) particles, such as ^56^Fe, and cannot be effectively blocked by modern spacecraft shielding (Cucinotta et al., 2006; Spillantini et al., 2007; Durante, 2014; Nelson, 2016; Zeitlin and La Tessa, 2016). Rodent data suggest HZE particle exposure is detrimental to brain physiology and functional cognitive output (Blakely and Chang, 2007; Cucinotta et al., 2014; Kokhan et al., 2016; Nelson, 2016; Jandial et al., 2018; Cucinotta and Cacao, 2019; Kiffer et al., 2019; Limoli, 2020; Davis et al., 2021). Taken together with the unavoidable nature of GCR, HZE particle exposure appears to pose an unavoidable threat to astronaut well-being and mission success. Specifically, a central theme emerging from rodent space radiation literature (which overwhelmingly has used ^56^Fe particles) is that exposure to HZE particles may be harmful to astronaut cognition and brain health.

A thorough review of the literature, however, does not support a uniformly negative impact of HZE particles on rodent brain and behavior (Kiffer et al., 2019). Two even more recent studies highlight that ^56^Fe particle exposure can have seemingly beneficial effects on the mouse hippocampus, a brain region intimately involved with memory and mood regulation. One study showed exposure to whole body ^56^Fe particle irradiation (IRR) improves hippocampal-dependent spatial learning 12 and 20 mon post-radiation in male and female mice (Miry et al., 2021). Another study exposed 6-mon old (“astronaut-age”) male mice to whole body ^56^Fe particles (Whoolery et al., 2020) and used a rodent touchscreen platform to probe the functional integrity of brain circuits (Oomen et al., 2013; Hvoslef-Eide et al., 2016; Kangas and Bergman, 2017), drawing similarity to the way astronauts undergo touchscreen testing (Basner et al., 2017; Moore et al., 2017). This study found mice exposed to ^56^Fe particles had better discrimination learning (location discrimination, LD) vs. Sham mice, suggesting astronauts may show an improvement in this mission-critical skill. However, ^56^Fe mice were not different from Sham mice in many other tasks (pairwise discrimination, PD; visuospatial/associative-learning, Paired Associates Learning, PAL; stimulus-response habit or “rule-based” learning, visuomotor conditional learning, VMCL; cognitive flexibility, PD reversal) (Whoolery et al., 2020). Taken together, these studies suggest caution in concluding that HZE particle exposure decreases rodent cognition; the reality is likely that there are time-, task-, species-dose-, energy-, etc. dependent effects (Miry et al., 2021). In addition, these studies point out the importance - recently underscored for the space radiation field (Britten et al., 2021) - of measuring multiple cognitive processes in rodents, thereby preventing NASA’s risk assessments from being based on a single cognitive domain.

Another important factor to consider in assessing the impact of HZE particle exposure on cognition is biological sex. As of 2019, <10% of preclinical studies assessing the cognitive effects of HZE particle exposure use female rodents (Kiffer et al., 2019). Human research is only slightly better; the available data from astronauts is heavily skewed in favor of males (n = 477) vs. females (n = 57). Thus there is not enough data available to understand what role biological sex plays in the body’s response to space flight stressors (Mark et al., 2014), including exposure to GCR and HZE particles. It is intriguing, though, that some preclinical work suggests the female rodent brain may be protected from radiation-induced immune and cognitive deficits (Villasana et al., 2011; Krukowski et al., 2018), or at least that there are sex-specific effects in the impact of ^56^Fe exposure on hippocampal-dependent function (Villasana et al., 2010, 2011). Considering the astronaut class is 40% women (Mark et al., 2014), and the translational relevance of the rodent touchscreen platform (Oomen et al., 2013; Hvoslef-Eide et al., 2016; Kangas and Bergman, 2017), it is striking that the touchscreen platform has not yet been used to assess how female rodent cognition is influenced by whole-body exposure to an HZE particle, such as ^56^Fe.

To address this knowledge gap, mature C57BL/6J female mice received either Sham or whole-body ^56^Fe particle irradiation (IRR; 3 × 6.7cGy ^56^Fe, 600MeV/n) and were assessed on a battery of touchscreen and classical behavior tasks to assess aspects of cognition. These radiation exposure parameters are identical to those that reportedly improve location discrimination in mature male mice (Whoolery et al., 2020), and this dose was chosen as it is submaximal to that predicted for a Mars mission (Cucinotta and Durante, 2006; Hellweg and Baumstark-Khan, 2007). For the present study, female Sham and IRR mice were tested for touchscreen performance of instrumental learning, discrimination learning, extinction learning, and stimulus-response habit (rule-based) learning. Given literature suggesting the female rodent brain may be spared from the negative impact of HZE particle exposure (Rabin et al., 2013; Krukowski et al., 2018), we hypothesized that whole-body ^56^Fe IRR would spare or even improve their performance in touchscreen-based behaviors, especially hippocampal-reliant discrimination learning. This touchscreen battery revealed an unexpected finding: improved discrimination learning, but worse stimulus-response habit learning in ^56^Fe-irradiated vs.Sham mice. We also tested several classical behaviors to examine these anxiety, stress response and repetitive behaviors, but both Sham and IRR mice performed similarly in those tests. Taken together with pre-planned, in-depth analysis of key aspects of their touchscreen performance on this and many other tasks, these data suggest whole-body exposure to ^56^Fe particle IRR in mature female mice may support or enhance hippocampal tasks like discrimination learning, but may diminish to striatum-dependent tasks like stimulus-response habit learning.

## 2. Material and Methods

The ARRIVE 2.0 guidelines were used to design and report this study (Percie du Sert et al., 2020). A protocol was prepared for this study prior to experimentation, but this protocol was not registered.

### 2.1. Animals

Two-mon-old female C57BL/6J mice were purchased from Jackson Laboratories (stock #000664) and housed at Children’s Hospital of Philadelphia (CHOP, **Fig. 1A**) or UT Southwestern Medical Center (UTSW, **Fig. 1B**) and shipped to Brookhaven National Laboratories (BNL) for irradiation at 6-mon of age. Housing conditions at all facilities are 3-4/cage, light on 06:00, lights off 18:00, UTSW/CHOP: room temperature 68-79°F, room humidity 30-70%, BNL: room temperature 70-74°F and room humidity 30-70%. During shipping and housing at BNL, mice were provided Shepherd Shacks (Bio-Serv); no other enrichment was provided during housing. After IRR, mice were transferred to either UTSW or CHOP. At both facilities, food (CHOP and BNL: LabDiet #5015; UTSW: Envigo Teklad global 16% protein) and water were provided *ad libitum* except during the appetitive behavior tasks. When placed in CHOP quarantine after return from IRR (below), all mice received modified chow (Test Diet, Cat#1813527, Modified LabDiet 5058 with 13ppm Ivermectin and 150 ppm Fenbendazole) as required by CHOP’s Department of Veterinary Research. Animal procedures and husbandry were in accordance with the National Institutes of Health Guide for the Care and Use of Laboratory Animals, and performed in IACUC-approved facilities at UTSW (Dallas TX; AAALAC Accreditation #000673, PHS Animal Welfare Assurance D16-00296, Office of Laboratory Animal Welfare [OLAW] A3472-01), CHOP (Philadelphia, PA; AAALAC Accreditation #000427, PHS Animal Welfare Assurance D16-00280 [OLAW A3442-01]) and BNL (Upton NY; AAALAC Accreditation #000048, PHS Animal Welfare Assurance D16-00067 [OLAW A3106-01]). Of the 52 total mice used for this study, 3 mice (n=2 Sham, n=1 IRR) had to be euthanized for reaching a humane endpoint (lethargy, hunched posture, coat unkempt). None of the other mice used in this study warranted us employing our established interventions for reducing pain, suffering, or distress.

**Figure 1.**
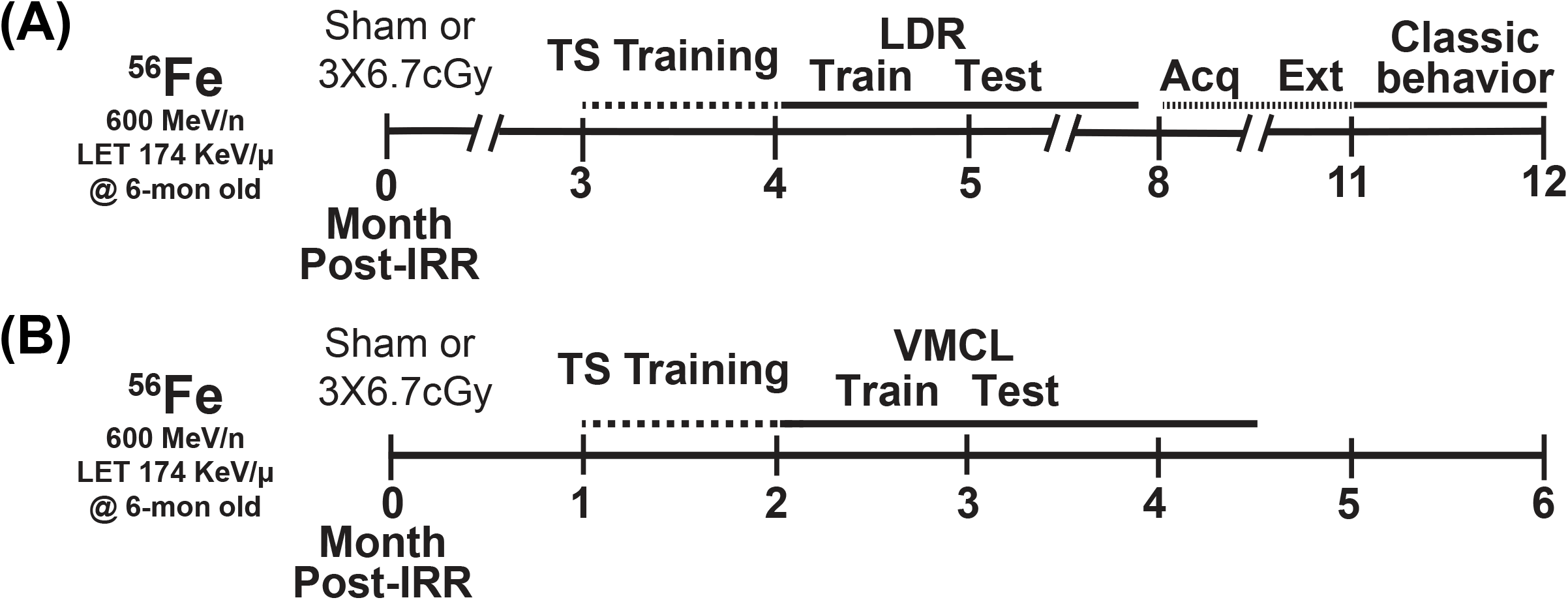
Timeline of experimental groups and overview of behavior tests. Six-month old C57BL/6J female mice received whole-body exposure to ^56^Fe (0-Month Post-Irradiation [IRR]) and subsequently were run on a variety of touchscreen and non-touchscreen behavioral tests, including (**A**)touchscreen training with a twelve-window (2 × 6) grid followed by LDR Train and Test, Acquisition and Extinction of stimulus-response habit learning, and classic behavior tests (EPM, MB, OF, SI, FST) or (**B**) touchscreen training on a 3-window (1 × 3) grid followed by VMCL Train and Test. Acq, acquisition; Ext, extinction; LDR, location discrimination reversal; Mon, month; Test, testing; Train, training; TS, touchscreen; VMCL, visuomotor conditioning learning.

### 2.2. Particle Irradiation (IRR)

Mice received whole-body HZE particle IRR at BNL’s NASA Space Radiation Laboratory (NSRL) during NSRL campaigns 17B and 18A. The ^56^Fe ion beams were produced by the AGS Booster Accelerator at BNL and transferred to the experimental beam line in the NSRL. Dosimetry and beam uniformity was provided by NSRL staff. Delivered doses were ±0.5% of the requested value. All mice - regardless of whether control (Sham) or experimental (^56^Fe) - were singly-placed for 15 minutes (min) in modified clear polystyrene rectangular containers (AMAC Plastics, Cat #100C, W 5.8 × L 5.8 × H 10.7 cm; modified with ten 5-mm air holes). Although confined to a container, mice had room to move freely and turn around during confinement. A maximum of six containers were placed perpendicular to the beam for each cave entry. Mice received either Sham exposure (placed in cubes Monday, Wednesday, Friday, but received no ^56^Fe exposure) or Fractionated (Frac) 20 cGy ^56^Fe IRR (600 MeV/n, LET 174 KeV/μ, dose rate 20 cGy/min; placed in cubes and received 6.7 cGy on Monday, Wednesday, and Friday). Post-IRR, mice were returned to UTSW (**Fig. 1B**) or CHOP (**Fig. 1A**) and housed in quarantine for 1-1.5 mon prior to initiation of touchscreen behavior testing (**Fig. 1**). Body weights (**Fig. 2A**) were taken multiple times: prior to IRR, at IRR, and at least weekly post-IRR until collection of brain tissue. This dose of ^56^Fe was selected as it is submaximal to that predicted for a Mars mission (Cucinotta and Durante, 2006; Hellweg and Baumstark-Khan, 2007) and the fractionation interval (48 hours [hr]) was determined by the inter-fraction period for potential repair processes (Thames, 1985).

**Figure 2.**
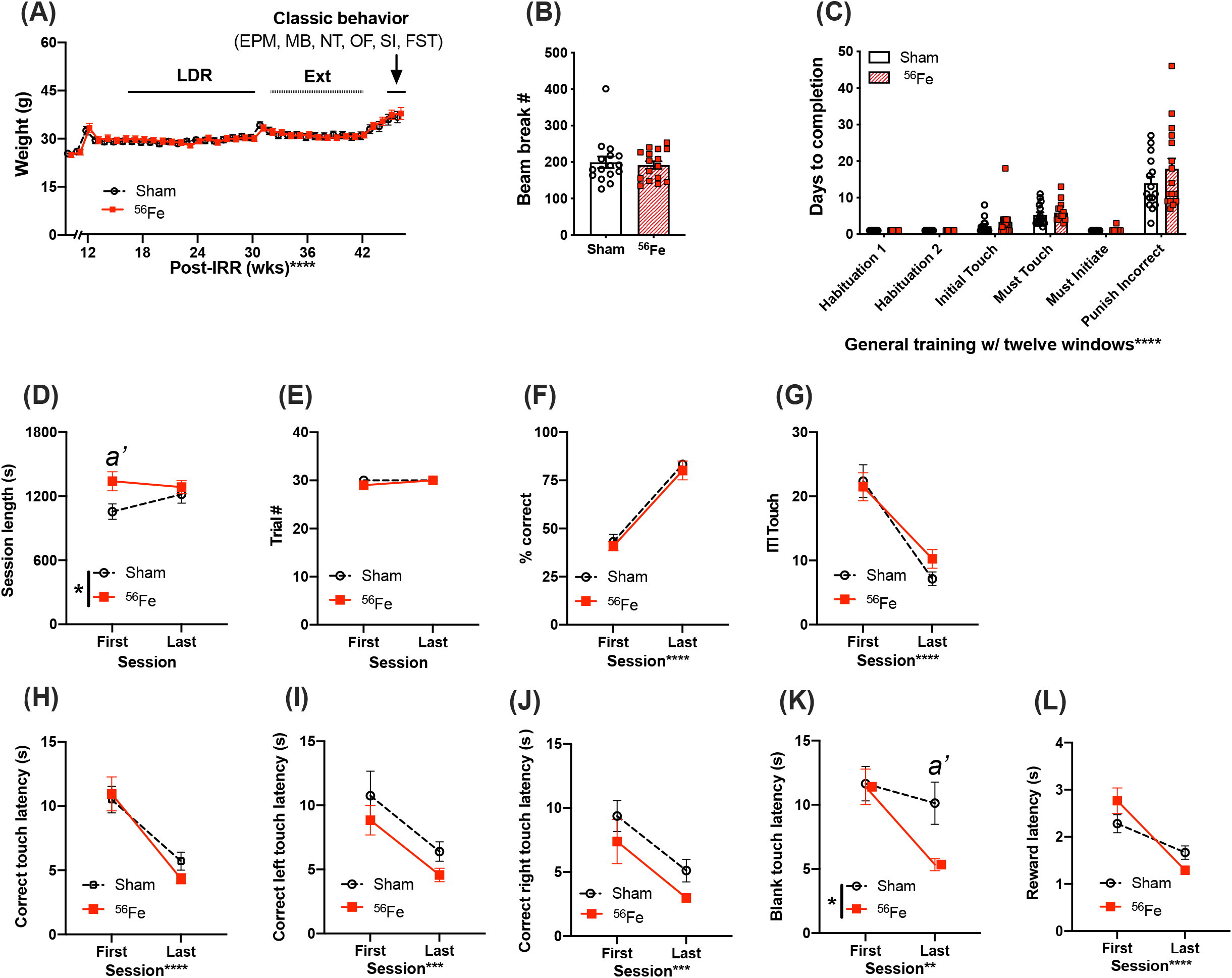
Weights, locomotion, and general touchscreen training learning are generally unaffected in 6-mon old female mice exposed to whole-body ^56^Fe IRR compared to Sham. **(A)** No gross weekly weight difference was detected between Sham or ^56^Fe mice during touchscreen testing. **(B)** Beam breaks measured in the novel TS operant chambers in Habituation 1 revealed no gross baseline difference in locomotion after exposure to Sham or ^56^Fe IRR. **(C)** Sham and ^56^Fe IRR groups performed similarly in each of the first six steps of general touchscreen training with twelve windows: Habituation 1 and 2, Initial Touch, Must Touch, Must Initiate, and Punish Incorrect. **(D-L)** During the Punish Incorrect stage of general TS training, ^56^Fe IRR female mice had a longer first, but not last, training session vs. Sham mice **(D)**. However, Sham and ^56^Fe IRR mice did not differ in the number of completed trials **(E)**, % correct **(F)**, total ITI touches **(G)**, correct touch latency **(H)**, and correct left or right touch latency **(H, I). (K)** ^56^Fe IRR female mice were ∼5s faster vs. Sham mice to touch a blank window in the final session of testing. Aside from these differences, IRR and Sham mice had similar reward collection latency **(L)**. Error bars depict mean±SEM. Statistical analysis in **A, C-L**: Mixed-effects analysis **(A)**, Two-way RM ANOVA **(C-L)**, main effect *p<0.05, **p<0.01, ***p<0.001, ****p<0.0001, Bonferroni’s post-hoc analysis *a’*<0.05 and in **B**: Unpaired t-tests. EPM, elevated plus maze; FST, forced swim test; IRR, irradiation; MB, marble burying; OF, open field; SI, social interaction; TS, touchscreen; s, seconds; wks, weeks.

### 2.3. Overview of behavioral testing

Mice exposed to HZE particles in NSRL campaigns 18A and 17B were divided into parallel groups (**Fig. 1A, B**, respectively) that underwent touchscreen behavioral testing 1-3 mon post-IRR. Touchscreen experiments were performed between 8:00am to 2:00pm during weekdays. As is standard in most rodent touchscreen experiments, mice were food restricted during touchscreen experiments. Mouse chow was removed from each cage at 5 pm the day prior to training or testing. Each cage was given *ad libitum* access to chow for 3 hour (hr; minimum) to 4 hr (maximum) immediately following daily touchscreen training/testing, and from completion of training/testing on Friday until Sunday 5 pm. Mice were weighed each Wednesday to ensure weights >80% initial body weight. While weights below this threshold merited removal of the mouse from the study, zero mice reached this threshold (Mar et al., 2013; Oomen et al., 2013). Luminescence emits from the touchscreen chamber screen and reward magazine, and from the house light during one stage of general touchscreen training, and thus the mice are not performing in darkness. In one group of mice (**Fig. 1A**), mice began touchscreen behavioral testing 3-mon post-IRR. Operant touchscreen platform procedures included general touchscreen training (with 2 × 6 window grid), Location Discrimination Reversal [LDR, training and testing], and Extinction [Ext, training or “Acqusition” and testing]. Total beam breaks as a measure of baseline locomotor activity were gathered in touchscreen operant chambers during general touchscreen training (Habituation 1). After all animals completed LDR testing, mice received unrestricted food pellets for two weeks before beginning Ext to allow them to recover from potential stress associated with food restriction. Following Ext testing, mice were tested in a variety of non-touchscreen tests classified here as “classical behavior tests” (**Fig. 1A, 2A, 6**).

These tests measure anxiety-(elevated plus maze [EPM], open field [OF]) or repetitive/compulsive-like behaviors (marble burying [MB]), locomotion (OF), sociability (social interaction [SI]), and despair-like behaviors (forced swim test [FST]), methods for which are provided below. Classical behavior tests were performed between 2:00pm to 5:00pm during weekdays under red light (45-65 lux) except for the forced swim test which was done under white and red light simultaneously. Mice were habituated to the testing suite under red light for 1h prior to testing. In the second group of mice (**Fig. 1B**), mice began touchscreen behavioral testing 1-mon post-IRR. Operant touchscreen platform procedures performed on this group were general touchscreen training [with 1 × 3 window grid] and Visuomotor Conditional Learning [VMCL] training and testing.

#### 2.3.1. General Touchscreen Training (prior to LDR)

consists of 6 stages, as previously published (Whoolery et al., 2020): Habituation 1, Habituation 2, Initial Touch, Must Touch, Must Initiate, and Punish Incorrect (PI). Methods for each stage are described in turn below. Mice went through general touchscreen training with twelve windows (2 × 6) for the LDR experiment.

##### 2.3.1.1. Habituation

Mice are individually placed in a touchscreen chamber for 30-min (max) with the magazine light turned on (LED Light, 75.2 lux). For the initial reward in each habituation session, a tone is played (70 decibel [dB] at 500 Hz, 1000 ms) at the same time as a priming reward (150-ul Ensure® Original Strawberry Nutrition Shake) is dispensed to the reward magazine. After a mouse inserts and removes her head from the magazine, the magazine light turns off and a 10-s delay begins. At the end of the delay, the magazine light is turned on and the tone is played again as a standard amount of the reward (7-ul Ensure) is dispensed. If the mouse’s head remains in the magazine at the end of the 10-s delay, an additional 1-s delay is added. A mouse completes Habituation training after they collect 25 rewards (25 × 7 ul) within 30 min. Mice that achieve habituation criteria in <30 min are removed from the chamber immediately after their 25th reward in order to minimize extinction learning. The measure reported for Habituation is days to completion.

##### 2.3.1.2. Initial Touch

A 2 × 6 window grid is placed in front of the touchscreen for the remaining stages of training. At the start of the session, an image (a lit white square) appears in a pseudo-random location in one of the 12 windows on the touchscreen. The mouse has 30 s to touch the lit square (typically with their nose). If the mouse does not touch the image, it is removed, a reward (7ul Ensure) is delivered into the illuminated magazine on the opposite wall from the touchscreen, and a tone is played. After the reward is collected, the magazine light turns off and a 20-s intertrial interval (ITI) begins. If the mouse touches the image while it is displayed, the image is removed and the mouse receives 3 times the normal reward (21-ul Ensure, magazine is illuminated, tone is played). For subsequent trials, the image appears in another of the 12 windows on the touchscreen, and never in the same location more than 3 consecutive times. Mice reach criteria and advance past Initial Touch training when they complete 25 trials (irrespective of reward level received) within 30 min. Mice that achieve Initial Touch criteria in <30 min are removed from the chamber immediately after their 25th trial. The measure reported for Initial Touch is days to completion.

##### 2.3.1.3. Must Touch

Similar to Initial Touch training, an image appears, but now the window remains lit until it is touched. If the mouse touches the lit square, the mouse receives a reward (7-ul Ensure, magazine is illuminated, tone is played). If the mouse touches one of the blank windows, there is no response (no reward is dispensed, the magazine is not illuminated, and no tone is played). Mice reach criteria and advance past Must Touch training after they complete 25 trials within 30 min. Mice that achieve Must Touch criteria in <30 min are removed from the chamber immediately after their 25th trial. The measure reported for Must Touch is days to completion.

##### 2.3.1.3. Must Initiate

Must Initiate training is similar to Must Touch training, but a mouse is now required to initiate the training by placing its head into the already-illuminated magazine. A random placement of the image (lit white square) will then appear on the screen, and the mouse must touch the image to receive a reward (7-ul Ensure, magazine lit, tone played). Following the collection of the reward, the mouse must remove its head from the magazine and then reinsert its head to initiate the next trial. Mice advance from Must Initiate training after they complete 25 trials within 30 min. Mice that achieve Must Initiate criteria in <30 min are removed from the chamber immediately after their 25th trial. The measure reported for Must Initiate is days to completion.

##### 2.3.1.4. Punish Incorrect (PI)

Punish Incorrect training builds on Must Initiate training, but here if a mouse touches a portion of the screen that is blank (does not have a lit white square), the overhead house light turns on and the lit white square disappears from the screen. After a 5-s timeout period, the house light turns off and the mouse has to initiate a correction trial where the lit white square appears in the same location on the screen. The correction trials are repeated until the mouse successfully presses the lit white square; however, correction trials are not counted towards the final percent correct criteria. Mice reach criteria and advance past Punish Incorrect training and onto Location Discrimination Reversal Train/Test after they complete 30 trials within 30 min at ≥ 76% (≥ 19 correct) on day 1 and >80% (>24 correct) on day 2 over two consecutive days. Mice that achieve Punish Incorrect criteria in <30 min are removed from the chamber immediately after their 30th trial. As with the other stages, a measure reported for Punish Incorrect is days to completion (to reach criteria). However, since the Punish Incorrect stage also contains a metric of accuracy, more measures were analyzed relative to the other five stages. Therefore, other measures reported for Punish Incorrect are session length, trial number, percent correct responses, intertrial interval (ITI), latency to make a correct touch (for total touches, left touches, and right touches) and an incorrect touch (touching a blank window), and latency to collect a reward.

#### 2.3.2 Location Discrimination Reversal

(LDR; program LD1 choice reversal v3; ABET II software, Cat #89546-6) tests the ability to discriminate two conditioned stimuli that are separated either by a large or small separation. The reversal component of LDR is used here and in classic LDR studies (Clelland et al., 2009; Oomen et al., 2013) tests cognitive flexibility; prior work showing space radiation improved LD function in male mice used LD, not LDR (Whoolery et al., 2020). Taken together, LDR is a hippocampal-dependent task (Clelland et al., 2009; Oomen et al., 2013) which allows assessment of both discrimination ability as well as cognitive flexibility. In our timeline **(Fig. 1A)**, mice received one additional training step (“LDR train”) prior to the actual 2-choice LDR test.

##### 2.3.2.1. Location Discrimination Reversal Train (LDR Train)

Mice initiated the trial, which led to the display of two identical white squares (25 × 25 pixels, **Fig. 3A**) presented with two blank (unlit) squares between them, a separation which was termed “intermediate” (8th and 11th windows in 2 × 6 high grid-bottom row). One of the left (L) or right (R) locations of the squares was rewarded (i.e. L+) and the other is not (R-), and the initial rewarded location (left or right) was counterbalanced within-group. On subsequent days, the rewarded square location was switched based on the previous day’s performance (L+ becomes L- and R-becomes R+, then L-becomes L+ and R+ becomes R-, etc). A daily LDR train session is complete once the mouse touches either L+ or R-50 times or when 30 min has passed. Once 7 out of 8 trials had been correctly responded to, on a rolling basis, the rewarded square location was switched (becomes L-), then L+, then L-, etc.; this is termed a “reversal”. Once the mouse reached >1 reversal in 3 out of 4 consecutive testing sessions, the mouse advanced to the LDR Test. A daily training is considered a “session”. Measures reported for LDR Train are: percent of each group reaching criteria over time (survival curve), days to completion, trial number, and percent correct during trials to the 1st reversal,

**Figure 3.**
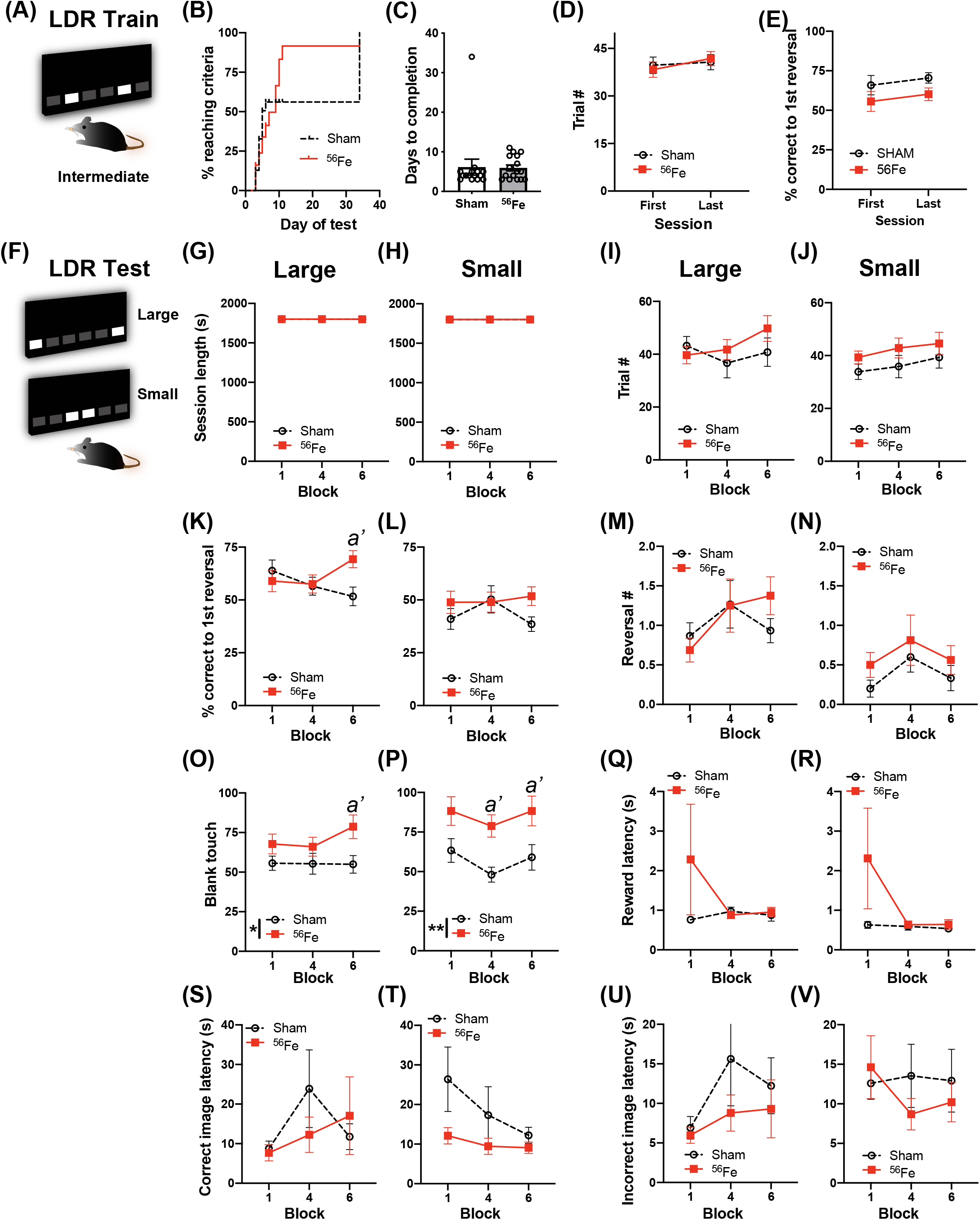
On an appetitive, touchscreen discrimination learning task, female mice exposed to whole-body ^56^Fe IRR at 6-mon of age perform better than Sham mice in discriminating the location of two identical visual cues. **(A)** Schematic of the lit squares and their Intermediate separation which are used for location discrimination reversal training (LDR Train). **(B-E)** Sham and ^56^Fe IRR mice performed similarly in LDR Train based on **(B)** distribution of subjects reaching criteria (the visual difference in percent of Sham and ^56^Fe subjects reaching criteria was rejected by survival curve analysis), **(C)** days to completion, **(D)** trials completed, and **(E)** % correct. **(F)** Schematic of the lit squares and their Large and Small separation used for LDR testing (LDR Test). **(G-X)** When mice underwent LDR Test with squares maximally-separated (Large separation, **G**,**I**,**K**,**M**,**O**,**Q**,**S**,**U**,**W**) or minimally-separated (Small separation, **H**,**J**,**L**,**N**,**P**,**R**,**T**,**V**,**X**), there was no difference between IRR and Sham mice in session length **(G**,**H)** or number of completed trials **(I**,**J)** on the last day of the 1st, 4th, and 6th two-day block. However, in the Large - but not Small - separation trials, ^56^Fe IRR female mice had greater accuracy vs. Sham mice in trials to the first reversal **(K**,**L)**. Sham and IRR groups did not differ in the number of reversals completed **(M**,**N)**. Both Large and Small separation trials revealed that ^56^Fe IRR female mice made more blank touches (to non-stimuli windows) vs. Sham mice **(O**,**P)**, specifically during the 6th (last) block of Large separation trials and during the 4th and 6th blocks of Small separation trials. IRR and Sham mice had similar reward collection latency **(Q**,**R)**, correct image response latency **(S**,**T)**, and incorrect image response latency **(U**,**V)** in the Large and Small separation trials. Error bars depict mean±SEM. Statistical analysis in **D-V**: Two-way RM ANOVA, main effect *p<0.05, **p<0.01, Bonferroni’s post-hoc analysis *a’*<0.05, in **B:** Mantel-Cox test, and **C:** Unpaired, two-tailed t-tests. LDR, location discrimination reversal; s, seconds.

##### 2.3.2.2. Location Discrimination Reversal Test (LDR Test)

Mice initiated the trial, which led to the display of two identical white squares, either with four black squares between them (“large” separation, two at maximum separation [7th and 12th windows in the bottom row of a 2 × 6 grid] or directly next to each other (“small” separation, two at minimum separation [9th and 10th windows in the bottom row of a 2 × 6 grid; **Fig. 3F**]). As in LDR Train, only one of the square locations (right-most or left-most) was rewarded (L+, same side for both large and small separation, and counterbalanced within-groups). The rewarded square location was reversed based on the previous day’s performance (L+ becomes L-, then L+, then L-, etc). Once 7 out of 8 trials had been correctly responded to, on a rolling basis, the rewarded square location was reversed (becomes L-, then L+, then L-, etc.). Each mouse was exposed to only one separation type during a daily LDR Test session (either large or small) and the separation type changed every two days (two days of large, then two days of small, two days of large, etc.). A daily LDR Test session was completed once the mouse touched either L+ or R-81 times or when 30 min had passed. LDR Test data are analyzed by block (1 block = 4 days LDR Test counterbalanced with 2 Large and 2 Small separation daily sessions). Once 24 testing sessions (12 days of Large, 12 days of Small separation) were completed, mice received a two-week normal feeding prior to extinction testing. Measures reported for LDR Test are all presented for both Large and Small separation: session length, trial number, percent correct during trials to the 1st reversal, number of reversal, number of blank touches (touching an un-lit square), reward collection latency, latency to touch the correct image on the last day of the 1st, 4th, and 6th two-day block were reported, and latency to touch the incorrect image (touching the incorrect lit square; does not include blank window touches) on the last day of the 1st, 4th, and 6th two-day block (to allow assessment in the last day in Large or Small separation testing blocks).

#### 2.3.3. Extinction learning (Ext; ABET II software, Cat #89547)(Mar et al., 2013)

##### 2.3.3.1. Acquisition of simple stimulus-response learning (schedule name: Extinction pt 1)

is the first part of the extinction test, and is a task that involves the amygdala (Fernando et al., 2013). The start of acquisition was marked by the magazine light turning on and the delivery of a free reward. Mice initiate the trial, which leads to the display of an image (lit white square stimulus) in the center window (middle square in 1 × 3 grid; **Fig. 4A**). The mouse must touch the stimulus displayed in the center window to elicit tone/food response. The two side windows were left blank throughout the experiment. No response ensued if the mouse touches a blank part of the screen. A daily acquisition session was complete once the mouse touched the center window 30 times or when 30 min had passed. Once the mouse completed 30 trials within 15 min on each of five consecutive sessions (criteria for acquisition), the mouse advanced to the extinction test. Measures reported for Acquisition are: percent of each group reaching criteria over time (survival curve), days to completion, session length, and number of correct responses.

**Figure 4.**
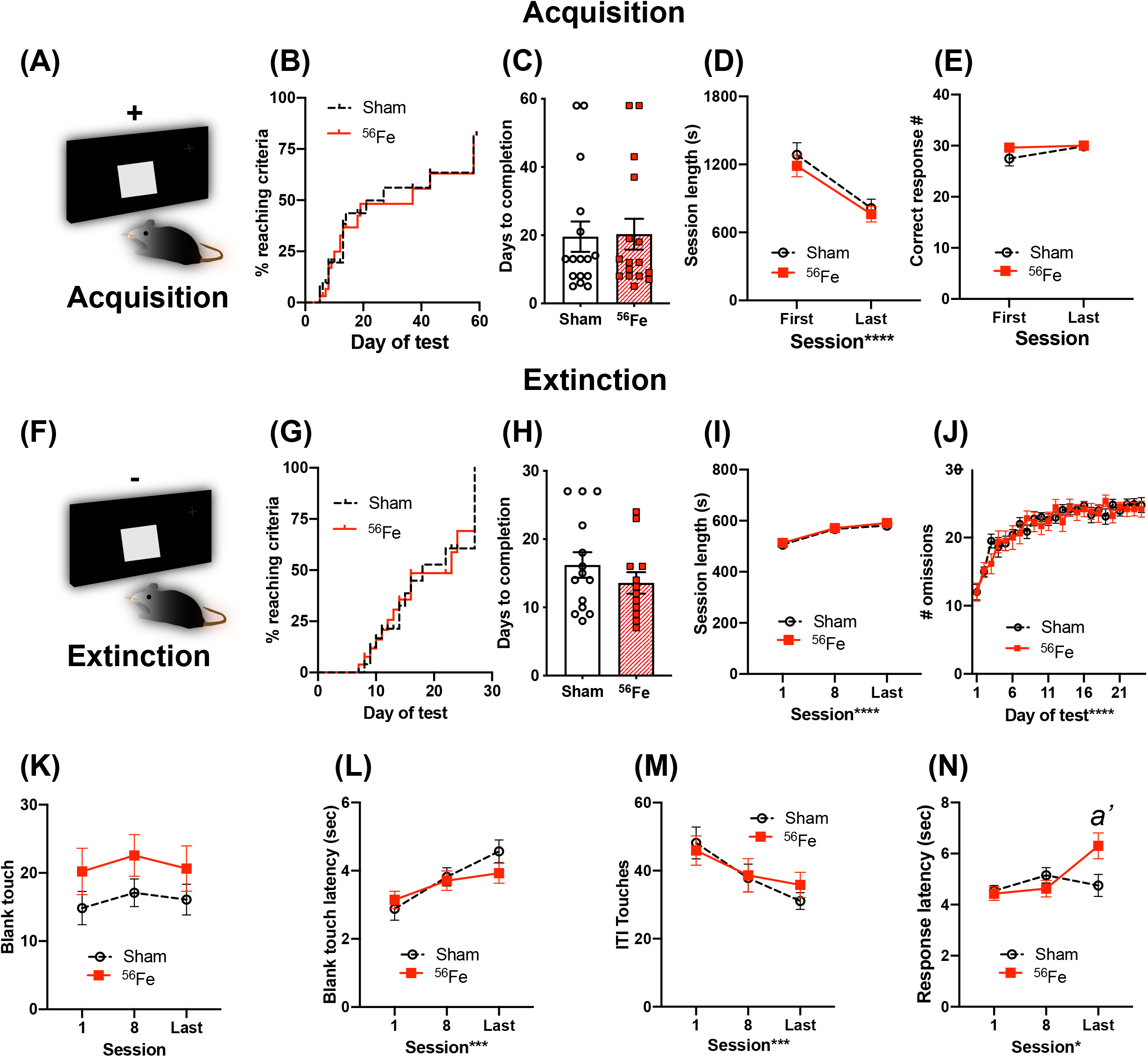
6-mon-old female mice exposed to whole-body ^56^Fe IRR and Sham mice perform similarly in acquisition and extinction of simple stimulus-response operant tasks. **(A)** Schematic of the single, large lit square image used for Acquisition of stimulus-response operant learning. Plus (+) sign indicates that the mouse is rewarded for touching the single, large, lit square. **(B-E)** Sham and ^56^Fe IRR mice performed similarly during stimulus-response Acquisition, based on distribution of subjects reaching criteria, **(C)** days to completion, **(D)** session length, and **(E)** number of correct responses. **(F)** Schematic of the single, large lit square image used for extinction of the operant stimulus-response learning. Minus (-) sign indicates that the mouse receives no reward during extinction. **(H)** ^56^Fe IRR and Sham mice took similar time to extinguish the previously-acquired stimulus-response contingent behavior, and the performance of Sham vs. IRR mice was similar based on session length **(I)**, number of omissions **(J)**, number of blank touches **(K)**, blank touch latency **(L)**, and ITI touches **(M)**. ^56^Fe IRR mice showed a higher response touch latency than Sham mice in the last session of extinction testing **(N)**. Error bars depict mean±SEM. Statistical analysis in **D**,**E**,**I**,**K-N**: Two-way RM ANOVA, main effect *p<0.05, **p<0.01, ****p<0.0001. Bonferroni’s post-hoc analysis *a’*<0.05, in **B**,**G:** Mantel-Cox test, and in **C**,**H**: Unpaired, two-tailed t-tests, *p<0.05. s, seconds.

##### 2.3.3.2. Extinction test (schedule name: Extinction pt 2)

is a test that involves the prefrontal cortex (Mar et al., 2013). A 5-s ITI marked the start of extinction. Following the initial ITI, an image (the lit white square stimulus) was presented in the center window (middle window in 1 × 3 grid; **Fig. 4F**). The stimulus display was held on the screen for 10 s during which the mouse could elicit or omit its learned response to the square. The two side windows were left blank throughout the experiment. If the mouse touched the blank window of the screen, no response occurred. If the white square was touched, no food delivery was made but the image was removed, the magazine light was illuminated, a tone was played and the ITI period (10 s) was started; this is the “correct” action, even though no reward is provided. If the white square was not touched, then the image was removed and the ITI period started. Mouse entry into the reward magazine during the ITI would turn off the magazine light. Following an ITI, the magazine light was turned off and the next trial began automatically. A daily extinction session was complete once the mouse was presented with the white square stimulus 30 times. When the mouse reached ≥80% response omissions on each of at least three out of four consecutive sessions, the mouse was considered to have reached daily criteria. Mice that reached criteria first continued to be tested daily until all of the mice’s performance was synchronized and completed before advancing to classical behavior battery testing. Measures reported for Extinction are: percent of each group reaching criteria over time (survival curve), days to completion, and number of omissions across testing days as well as session length, number of touches and latency to touch to a blank part of the touchscreen, number of touches during the ITI, and latency to make a correct response on the last day of the first, 8th, and last testing session.

#### 2.3.4. Visuomotor Conditional Learning (VMCL, ABET software, Cat #89542)

is a stimulus-response habit (or rule-based) learning task reliant on the striatum (Horner et al., 2013; Delotterie et al., 2015).

##### 2.3.4.1. General Touchscreen Training (prior to VMCL)

occurred as in **2.3.1** with two differences: mice went through only one habituation stage (Habituation 2) and training occurred with a three window grid (1 × 3). As in **2.3.1**, the measure reported for these five general touchscreen stages is days to completion (to reach criteria). Due to computer issues, “days to completion” was the only metric extracted for this group, precluding in-depth accuracy analyses of Punish Incorrect as was done in the LDR cohort. After these five training stages, mice then went through VMCL Train and finally VMCL Test.

##### 2.3.4.2. VMCL Train (schedule name: Punish Incorrect II)

VMCL Train was designed to teach the mouse to touch two images (both lit white squares) on the screen in a specific order and in rapid succession. The first touch must be to an image presented in the center of the screen, and the second touch must be to an image presented either on the left or right of the screen. Specifically, after trial initiation, the mouse must touch a center white square (200×200 pixels; **Fig. 5A**), which then disappears after it is touched. A second white square immediately appears on either the left or right side of the screen in a pseudorandom style, such that a square was located on each side 5 out of 10 times, but not more than 3 times in a row. If the mouse selected the location with the second white square, a reward (7ul) was provided, and a 20-s ITI began. However, if the mouse selected the location without a lit white square, then the second stimulus was removed, the house light was illuminated for 5 s to indicate a timeout period, and then finally a 20-s ITI occurred. Then the mouse was presented with a correction trial which must be completed prior to a new set of locations being displayed. VMCL Train was complete when the mouse completed 2 out of 3 consecutive days of 25 trials in 30 min with ≥75% correct. Measures reported for VMCL Train are: days to completion and distribution of percent of mice which reach criteria over time (survival curve), as well as session length, trial number, % correct responses, and number of correction trials on the first and last VMCL Train session.

**Figure 5.**
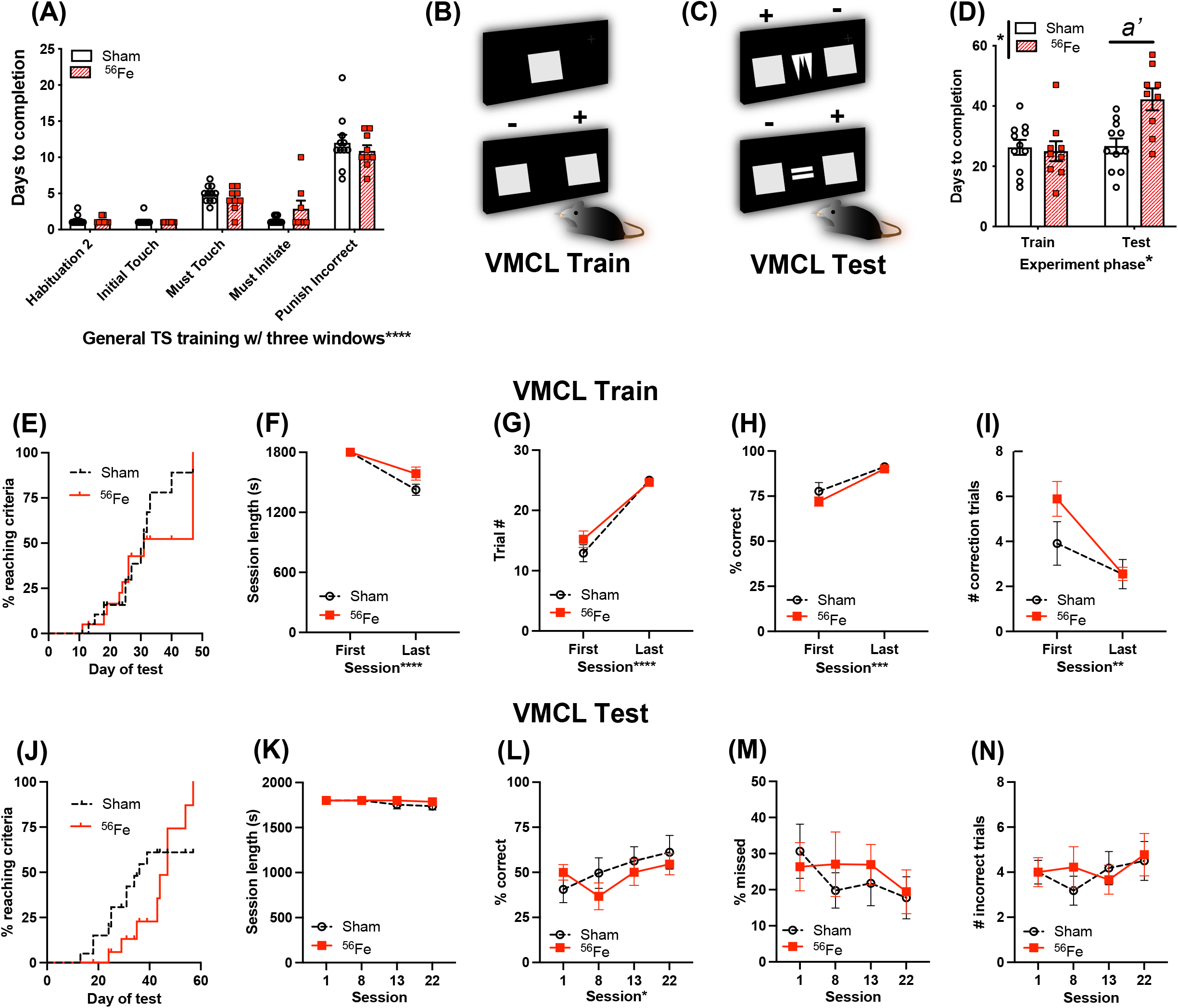
6-mon-old female mice exposed to whole-body ^56^Fe IRR perform worse than Sham in tests of stimulus-response habit learning. **(A)** Sham and ^56^Fe IRR groups performed similarly in each of the five steps of general touchscreen training with three windows: Habituation 2, Initial Touch, Must Touch, Must Initiate, and Punish Incorrect. **(B)** Schematic of the single, large lit square image (top) and two, large lit squares used for Visuomotor Conditional Learning (VMCL) Train. The mouse learns that touching one square (+) is rewarded while the other (-) is not rewarded. **(C)** Schematic of the two, large lit squares flanking samples used for VMCL Test. The mouse learns that touching one square (+) is rewarded while the other (-) is not rewarded depending on the center image shown. **(D)** ^56^Fe IRR mice took a similar number of days to complete VMCL Train - but took more days to complete the VMCL Test - compared to Sham mice. **(E)** Cumulative distribution function showed no difference between groups in days required to complete VMCL Train. **(F-I)** ^56^Fe IRR mice performed similarly as Sham mice in VMCL pre-testing (**F:** Session length, **G:** completed trials, **H:** % correct, **I:** # correction trials). **(J)** Cumulative distribution function showed no difference between groups in days required to complete VMCL Test. **(K-N)** All parameters (**K:** session length, **L:** % correct, **M:** % missed, **N:** #incorrect trials) between ^56^Fe IRR and Sham mice were similar. Error bars depict mean±SEM. Statistical analysis in **A**,**D**,**K-N**: Mixed-effect analysis, **F-I:** Two-way RM ANOVA, main effect *p<0.05, **p<0.01, ***p<0.001, ****p<0.0001. Bonferroni’s post-hoc analysis *a’*<0.05, and in **E**,**J:** Mantel-Cox test. s, seconds; TS, touchscreen; VMCL, Visuomotor Conditional Learning.

##### 2.3.4.3. VMCL Test

Mice were provided with one of two black-and-white images (spikes or horizontal bars, **Fig. 5C**) placed in the center window. Once touched, the center image disappeared and white squares simultaneously appeared on the right and left of the screen. For this task, the center image of the spikes signaled that the mouse should touch the right square, while the center image of the horizontal bars signaled that the mouse should touch the left square. The two center images were presented pseudorandomly for an equal number of times, and the mice had 2 s to touch the white square on the either right or left side of the central image depending on the center image type. If the mouse touched the appropriate image (right or left side), this was considered a correct trial, and the mouse received a reward (7ul) and then a 20-s ITI occurred. If the mouse touched the inappropriate image (right or left side), this was considered an incorrect trial, and the house light was illuminated for a 5-s timeout period followed by a 20-s ITI. If the mouse did not touch the white square (right or left side) within 2 s, this was considered a “missed” trial, and the house light was illuminated for a 5-s timeout period followed by a 20-s ITI. After either an incorrect or missed trial, a correction trial was runto protect against side bias. VMCL Test was completed when the mouse completed 2 consecutive days of 25 trials in 30 min with ≥ 80% correct responses. Measures reported for VMCL Test are: days to completion and distribution of percent of mice which reach criteria over time (survival curve), as well as session length, percent of responses that were correct, percent of missed responses, and number of incorrect trials on the 1st (first), 8th, 13th (intermediate), and 22nd (later) testing sessions (22^nd^ day chosen based on average days to meet the criteria of VMCL Test in Sham group).

#### 2.3.5. Classic Behavior Testing

##### 2.3.5.1 Elevated Plus Maze (EPM)

is a test for anxiety-like behavior (Yun et al., 2018). The maze contained two open arms and two closed arms each with a length of 67 cm and a width of 6 cm with the opaque walls of the closed arms being 17 cm tall (Harvard Apparatus, #760075). Mice were placed on the end of the open arms at the start of the behavior and allowed free movement throughout the maze for 5 min. The parameters of EPM (total distance of movement, entries and duration in the open arms, entries and duration in the closed arms) were scored via EthoVision software (Noldus Information Technology) using nose-center-tail tracking to determine position.

##### 2.3.5.2. Marble Burying (MB)

is a test of repetitive or compulsive-like behavior (Angoa-Pérez et al., 2013; Tran et al., 2020). Transparent polycarbonate cages (25.7 cm × 48.3 cm x15.2 cm) with filter-top lids (Allentown Inc. #PC10196HT) were used as marble burying arenas. 4-5 cm of wood chip bedding (Beta Chip Bedding, Animal Specialties and Provisions, #NOR301) was evenly distributed to cover the bottom of the cage and 20 glass marbles were laid gently on top of the bedding (four rows with five marbles in each row, evenly spaced). Mice were placed in the marble burying arena and given 20 minutes to explore and interact with the marbles. After 20 min of testing, marbles were scored by two independent observers and only marbles that were two-thirds or more covered in bedding were counted. The measure reported for this test is percent of marbles buried.

##### 2.3.5.3. Open Field (OF)

is a test for exploration and anxiety-like behavior (Yun et al., 2018; Tran et al., 2020). The open field arena measured 42 cm × 42 cm × 42 cm (opaque white Plexiglas, customer design Nationwide Plastics). The center zone was established in EthoVision as 14 cm × 14 cm and corner periphery zones were set as 5 cm × 5 cm each. Each mouse was placed in the arena and allowed free movement of the novel environment with recording for five minutes. The parameters of open field (total distance of movement, entries and duration in the center zone, entries and duration in the periphery zone) were scored via EthoVisionXT software (Noldus Information Technology) using nose-center-tail tracking to determine position.

##### 2.3.5.4. Social Interaction (SI)

is a test of exploration, locomotion, and sociability (Yun et al., 2018). Test mice were individually placed in a white open-field chamber (42 cm × 42 cm × 42 cm) that had a discrete interaction zone against one wall (26 cm × 14 cm) inside of which there was an empty plastic and wire mesh container (10 cm × 6 cm). For the first trial, the mouse was placed randomly into either corner of the box opposite to the interaction zone, and the movements of the mouse were tracked using Ethovision software (Noldus Information Technology). Specifically, the time the mouse spent either in the interaction zone or in corners opposite to the interaction zone during a 2.5 min trial was quantified. For the second trial, which began ∼5 min after the first trial, an unfamiliar age and sex-matched C57BL/6J mouse was placed into the plastic and wire mesh container and the container was placed in the interaction zone. Again, the time the mouse spent either in the interaction zone or in corners opposite to the interaction zone during a 2.5 min trial was quantified. Measures reported for Social Interaction are time spent in the interaction zone without and with another mouse placed inside the plastic and wire mesh container.

##### 2.3.5.5. Forced Swim Test (FST)

is a test of despair-like responses (Yun et al., 2018). The FST was performed to evaluate behavioral withdrawal induced by stress. Mice were placed in a 5L beaker (Corning Inc. Life Sciences, Lowell, MA, USA) filled with 4L of 25±2°C water. The movements of the mouse were tracked using Ethovision software (Noldus Information Technology) for the entirety of the 6-min session. Mice each went through two, 6-min sessions on consecutive days. Immobile time was measured and the last 4-min of data are reported.

### 2.4. Rigor, Additional ARRIVE 2.0 Details, and Statistical Analysis

The experimental unit in this study is a single mouse. For behavioral studies, mice were randomly assigned to groups. Steps were taken at each experimental stage to minimize potential confounds. For example, mice from the two experimental groups (Sham and ^56^Fe IRR) were interspersed throughout housing racks at UTSW, CHOP, and BNL (to prevent effects of cage location) and were interdigitated for all weighing and behaviors (to prevent an order effect). Sample sizes were pre-determined via power analysis and confirmed on the basis of extensive laboratory experience and consultation with CHOP and PennMed statisticians as previously reported (Whoolery et al., 2020). Exact sample number for each group is provided in **Table S1**. Data for each group are reported as mean ± s.e.m. Testing of data assumptions (normal distribution, similar variation between control and experimental groups, etc.) and statistical analyses were performed in GraphPad Prism (ver. 9.0.0). All analyses were hypothesis-based and therefore pre-planned, unless otherwise noted in Results. Statistical approaches and results including statistical analysis significance (p-values) and effect size (when RM two-way ANOVA p<0.05: partial omega-squared ω_p_^2^ where ≤ 0.05 small, ≥ 0.06 medium, ≥ 0.14 large) are provided in Results and/or **Table S1**. Analyses with two groups were performed using an unpaired, two-tailed Student’s t-test. Analyses with more than two variables were performed using two-way ANOVA or Mixed-effects analysis with Bonferroni post hoc test; repeated measures (RM) were used where appropriate, as indicated in figure legends and **Table S1**. Analysis of the distribution of subjects reaching criteria between control and experimental groups (survival curve) was performed with the Mantel-Cox test and significance was defined as *p < 0.05. A total of n=8 mice (n=5 Sham, n=3 IRR) were outliers based on *a priori* established experimental reasons (n=1 Sham did not complete the “Punish Incorrect” stage even by Day 53; n=1 Sham and n=1 IRR did not complete “Acquisition” even by Day 58; n=3 Sham and n=2 ^56^Fe did not complete “Extinction”) and the data from these mice were excluded from LDR, Acquisition, and Extinction analyses, respectively. Due to health issues, n=2 Sham and n=1 IRR mice were not run on classical behavior tests. Experimenters were blinded to treatment until analysis was complete.

#### 2.4.1. Scripts

Prior to statistical analysis, extinction and acquisition data were sorted and extracted. We used a custom Python 3.8.3, SQLite3 2.6.0, and Tkinter 8.6 script developed by the Eisch Lab to extract, calculate needed values, and compile the data into a database. Extracting the data into an output CSV file was managed with another custom script, and these outputs were verified manually. Following this verification, the data were analyzed using GraphPad Prism 8 according to the tests detailed in the Statistical Analysis section. These scripts along with sample data files are available at https://github.com/EischLab/18AExtinction.

## 3. Results

### 3.1. Whole-body ^56^Fe IRR does not change body weight or locomotor activity and has modest effects on operant learning in female mice

Six-month-old female C57BL/6J mice received either Sham IRR or Frac whole-body 20 cGy ^56^Fe (3 exposures of 6.7 cGy every-other day, total 20 cGy; **Fig. 1**). This total dose is submaximal to that predicted for a Mars mission, and the fractionation interval (48 hours [hr]) was used to allow potential repair processes to occur (Thames, 1985; Cucinotta and Durante, 2006; Hellweg and Baumstark-Khan, 2007). Consistent with a prior report (Whoolery et al., 2020), this dose and fractionation interval of ^56^Fe do not interfere with normal weight gain between groups (**Fig. 2A; Table S1**, Mixed-effects analysis, main effects: Time F_36,1067_= 66.05, P < 0.0001 and Treatment F_1,30_= 0.02450, P = 0.8767; interaction: TreatmentXTime F_36,1067_= 1.586, P = 0.0161, post-hoc: all P > 0.05) or locomotor activity (**Fig. 2B; Table S1**, unpaired t-test, t_30_= 0.3919, P = 0.6979).

Beginning 3-mon post-IRR, Sham and ^56^Fe IRR female mice began training on a touchscreen platform extensively validated in rodents (**Fig. 1A**) (Horner et al., 2013; Oomen et al., 2013; Hvoslef-Eide et al., 2016). Mice initially went through six stages of general touchscreen testing (**Fig. 1A**), with performance reflecting operant learning. Sham and ^56^Fe IRR mice completed all stages of general touchscreen training in similar periods of time (**Fig. 2C; Table S1**, two-way RM ANOVA, main effects: Train stage F_5,145_= 59.27, P < 0.0001, ω_p_^2^ = 0.62 and Treatment F_1,29_ = 2.5111, P = 0.1239; interaction: Train stageXTreatment F_5,145_ = 1.035, P = 0.3992). Thus, there was no gross difference in operant performance between these Sham and ^56^Fe IRR mice.

The final stage of general touchscreen testing, Punish Incorrect (PI, where an incorrect trial results in timeout), was next analyzed to a greater extent for two reasons. First, PI is the sole general touchscreen training stage that has an accuracy criterion (% correct). Second, PI is the stage that takes the longest to learn. Thus by comparing first vs.last PI day and considering measures relevant to accuracy (including individual session length, trial #, % correct, correct touches and latency, intertrial interval (ITI) touches, blank window touches, and reward latency, **Fig. 2D-L**), we hypothesized we would gain greater insight into how ^56^Fe influenced operant learning (via changes in speed, impulsivity, motivation, side bias, etc.)(Mar et al., 2013; Swan et al., 2014). ^56^Fe IRR mice took longer than Sham mice to complete a maximum of 30 trials on the first -- but not the last -- PI session (**Fig. 2D; Table S1**, two-way RM ANOVA, main effect: Session F_1,29_= 0.5688, P = 0.4568; Treatment F_1,29_ = 4.532, P = 0.0419, post-hoc: *a’* P = 0.0228, in Sham vs. ^56^Fe, ω_p_^2^=0.07; interaction: SessionXTreatment F_1,29_ = 2.345, P = 0.1365). However, on the first and last PI day Sham and ^56^Fe IRR mice performed similarly in many other PI metrics, including number of trials completed (**Fig. 2E**, two-way RM ANOVA, main effects: Session F_1,29_= 3.742, P = 0.0629 and Treatment F_1,29_ = 3.742, P = 0.0629; interaction: SessionXTreatment F_1,29_ = 3.742, P = 0.0629), percent correct (**Fig. 2F; Table S1**, two-way RM ANOVA, main effects: Session F_1,29_ = 149.5, P < 0.0001, ω_p_^2^ = 0.69 and Treatment F_1,29_ = 0.5868, P = 0.4499; interaction: SessionXTreatment F_1,29_= 0.01728, P = 0.8963), ITI touches (**Fig. 2G; Table S1**; two-way RM ANOVA, main effects: Session F_1,29_= 49.58, P < 0.0001, ω_p_^2^ =0.44 and Treatment F_1,29_ = 0.3334, P = 0.5681; interaction: SessionXTreatment F_1,29_ = 1.138, P = 0.2949), latency of total correct touches to the left and right side of screen (**Fig. 2H; Table S1**, two-way RM ANOVA, main effects: Session F_1,29_= 43.99, P < 0.0001, ω_p_^2^=0.38 “large” and Treatment F_1,29_ = 0.2134, P = 0.6476; interaction: SessionXTreatment F_1,29_= 1.127, P = 0.2971), left correct touch latency (**Fig. 2I; Table S1**, two-way RM ANOVA, main effects: Session F_1,29_ = 16.07, P = 0.0004, ω_p_^2^=0.17 and Treatment F_1,29_ = 2.078, P = 0.1601; interaction: SessionXTreatment F_1,29_= 0.001474, P = 0.9696), and right correct touch latency (**Fig. 2J; Table S1**, two-way RM ANOVA, main effects: Session F_1,29_ = 15.34, P = 0.0005, ω_p_^2^=0.17 “large”, and Treatment F_1,29_ = 2.961, P = 0.0960; interaction: SessionXTreatment F_1,29_= 0.005288, P = 0.9425). In regard to touching a blank window when the stimulus was presented, ^56^Fe IRR mice had a nearly 5s shorter latency vs. Sham mice during the last session of PI (**Fig. 2K; Table S1**, two-way RM ANOVA, main effects: Session F_1,29_= 9.015, P < 0.0055, ω_p_^2^=0.12 and Treatment F_1,29_= 3.842, P = 0.05, post-hoc *a’* P = 0.0203 in Sham vs. ^56^Fe, ω_p_^2^=0.05; interaction: SessionXTreatment F_1,29_= 3.234, P = 0.0825), suggesting late-developing impulsivity in ^56^Fe IRR mice. Motivation did not appear changed, as Sham and ^56^Fe IRR mice had similar reward collection latencies (**Fig. 2L, Table S1**, two-way RM ANOVA, main effects: Session F_1,29_= 30.43, P < 0.0001, ω_p_^2^=0.34 and Treatment F_1,29_= 0.1001, P = 0.7540, and interaction SessionXTreatment F_1,29_= 5.175, P = 0.0305, post-hoc: all P > 0.05 in Sham vs. ^56^Fe, ω_p_^2^=0.07). This further analysis of PI training stage suggests that ^56^Fe irradiation has modest effects on certain aspects of operant learning (IRR mice are slower to finish in daily PI session and may be more impulsive late in PI), but in many other ways perform indistinguishably from Sham IRR mice in PI and all other operant learning stages.

### 3.2. Female mice given whole-body ^56^Fe IRR perform better than Sham IRR mice in an appetitive-based location discrimination reversal touchscreen task

As previously reported, male mice given whole-body ^56^Fe IRR perform better than Sham IRR mice in an appetitive-based location discrimination (LD) touchscreen task, suggesting improved behavioral discrimination or behavioral pattern separation (Whoolery et al., 2020). The effect of ^56^Fe IRR exposure on translationally-relevant female mouse touchscreen performance is unknown, and specifically its effect on discrimination and cognitive flexibility is unknown. These are important knowledge gaps, as the proportion of US astronauts that are female is increasing and some preclinical work shows the female rodent brain may be less susceptible after charged particle exposure vs.the male rodent brain (Krukowski et al., 2018; Parihar et al., 2020).

To test if whole-body ^56^Fe IRR improves discrimination learning in adult female mice as it does in adult male mice, and to probe if ^56^Fe IRR impacts cognitive flexibility, Sham and ^56^Fe female mice were also assessed on LDR performance, consisting of training and testing sessions (**Fig. 1**)(Whoolery et al., 2020). In the LDR Training sessions (LDR Train, **Fig. 3A-E**; presentation of a two-choice stimulus response with lit squares intermediately-separated), there was a visual difference in percent of Sham and ^56^Fe subjects reaching criteria, but this difference was rejected by survival curve analysis (**Fig. 3B, Table S1;** Log-rank (Mantel-Cox) test, P=0.8480). Sham and ^56^Fe also had similar average days to complete LDR Training (**Fig. 3C**, unpaired t-test, t_29_= 0.09399, P = 0.9258) and completed a similar number of LDR Train trials (**Fig. 3D; Table S1**, two-way RM ANOVA, main effects: Session F_1,29_= 1.549, P = 0.2233 and Treatment F_1,29_= 0.001294, P = 0.9716; interaction: SessionXTreatment F_1,29_= 0.4779, P = 0.4949) and also had similar accuracy before the first reversal (**Fig. 3E; Table S1**, two-way RM ANOVA, main effects: F_1,29_= 0.9060, P=0.3490 and Treatment F_1,29_= 3.713, P = 0.0638; interaction: SessionXTreatment F_1,29_= 8.180e-006, P = 0.9977), indicating similar competency during the overall LDR Train sessions.

Sham and ^56^Fe IRR female mice were then assessed on overall LDR performance (LDR test, **Fig. 3F-V**), considering performance when the LDR Test squares were maximally separated (Large separation, **Fig. 3G,I,K,M,O,Q,S,U; Table S1**) or minimally separated (Small separation, **Fig. 3H,J,L,N,P,R,T,V; Table S1**), and are analyzed by block (1 block = 4 days LDR counterbalanced with 2 Large and 2 Small separation daily sessions). Sham and ^56^Fe IRR took a similar amount of time to complete sessions for both the Large (**Fig. 3G; Table S1**, two-way RM ANOVA, main effects: Block F_2,58_= 2.002, P = 0.1443 and Treatment F_1,29_ = 0.002083, P = 0.9639; interaction: BlockXTreatment F_2,58_ = 0.002083, P = 0.9979) and Small separation LDR Test trials (**Fig. 3H; Table S1**, main effects: Block F_2,58_ = 2.002, P = 0.1443 and Treatment F_1,29_ = 0.002083, P = 0.9639; interaction: BlockXTreatment F_2,58_= 0.002083, P = 0.9979), completed a similar number of trials for both Large (**Fig. 3I; Table S1**, two-way RM ANOVA, main effects: Block F_2,58_ =1.737, P = 0.1851 and Treatment F_1,29_ = 0.4793, P = 0.4943; interaction: BlockXTreatment F_2,58_ =1.877, P = 0.1622) and Small separation LDR test trials (**Fig. 3J; Table S1**, two-way RM ANOVA, main effects: Block F_2,58_= 2.175, P = 0.1228 and Treatment F_1,29_= 1.916, P = 0.1768; interaction: BlockXTreatment F_2,58_ =0.07135, P = 0.931). Sham and ^56^Fe IRR mice performance was also assessed for location discrimination reversal learning, which provides insight into both discrimination learning and cognitive flexibility (Vivar et al., 2012; Swan et al., 2014; Graf et al., 2018). First, we assessed discrimination learning by analyzing the percent correct trials made in each group prior to the 1st reversal. In Large - but not Small - separation trials, ^56^Fe IRR female mice were ∼34% more accurate vs. Sham mice in percent correct trials prior to the 1st reversal (Large: **Fig. 3K; Table S1**, two-way RM ANOVA, main effects: Block F_2,58_= 0.5836, P = 0.5611 and Treatment F_1,29_= 1.230, P = 0.2765; interaction: BlockXTreatment F_2,58_= 3.761, P = 0.0291, post-hoc: a’ P = 0.0220 in Sham vs. ^56^Fe, ω_p_^2^=0.05; Small: **Fig. 3L; Table S1**, two-way RM ANOVA, main effect: Block F_2,58_= 0.5919, P = 0.5566 and Treatment F_1,29_ = 2.469, P = 0.1269; interaction: BlockXTreatment F_2,58_= 1.149, P = 0.3242). However, Sham and ^56^Fe IRR mice had a similar number of reversals (an index of cognitive flexibility) in both Large (**Fig. 3M; Table S1**, two-way RM ANOVA, main effects: Block F_2,58_ = 2.559, P = 0.0861 and Treatment F_1,29_= 0.1469, P = 0.7043; interaction: BlockXTreatment F_2,58_ = 1.034, P = 0.3619) and Small separation LDR Test trials (**Fig. 3N; Table S1**, two-way RM ANOVA, main effects: Block F_2,58_= 1.992, P = 0.1457 and Treatment F_1,29_= 1.777, P = 0.1928; interaction: BlockXTreatment F_2,58_ = 0.03173, P = 0.9688). Together these data suggest ^56^Fe IRR mice have better discrimination learning than Sham mice, but ^56^Fe IRR mice and Sham have similar cognitive flexibility.

We next probed how ^56^Fe IRR mice were achieving greater accuracy in the Large separation LDR Test sessions (via possible changes in speed, impulsivity, motivation, side bias, etc.)(Mar et al., 2013; Swan et al., 2014). ^56^Fe IRR female mice touched the blank, non-stimulus window more than Sham mice during the 6th (last) block of the Large separation LDR Test block (**Fig. 3O; Table S1**, two-way RM ANOVA, main effect: Block F_2,58_= 0.7784, P = 0.4639 and Treatment F_1,29_= 6.262, P = 0.0182, post-hoc: *a’* P = 0.0232, ω_p_^2^=0.09; interaction of BlockXTreatment F_2,58_= 0.9088, P = 0.4087) and the 4th and 6th blocks of the Small separation LDR Test block (**Fig. 3P; Table S1**, two-way RM ANOVA, main effects: Block F_2,58_= 2.185, P = 0.1216 and Treatment F_1,29_= 11.49, P = 0.0020, post hoc: *a’* P = 0.0204 at Block 4, *a’* P = 0.0292 at Block 6, ω_p_^2^=0.17; interaction: BlockXTreatment F_2,58_= 0.1156, P = 0.8910), implying IRR-induced increased impulsivity. ^56^Fe IRR and Sham mice had similar reward collection latencies in both the Large (**Fig. 3Q, Table S1**, two-way RM ANOVA, main effects: Block F_2,58_= 0.6710, P = 0.5151 and Treatment F_1,29_= 1.083, P = 0.3066; interaction: BlockXTreatment F_2,58_= 1.095, P = 0.3414) and Small separation LDR Test blocks (**Fig. 3R; Table S1**, two-way RM ANOVA, main effect: Block F_2,58_= 1.675, P = 0.1962 and Treatment F_1,29_= 1.971, P = 0.1709; interaction: BlockXTreatment F_2,58_ = 1.426, P = 0.2485). ^56^Fe IRR and Sham mice took just a long to press the correct selection on the display in the Large (**Fig. 3S**, two-way RM ANOVA, main effects: Block F_2,58_= 1.298, P = 0.2808 and Treatment F_1,29_= 0.2292, P = 0.6357; interaction: BlockXTreatment F_2,58_= 0.9595, P = 0.3891) and Small separation LDR Test blocks (**Fig. 3T; Table S1**, two-way RM ANOVA, main effect: Block F_2,58_ = 1.979, P = 0.1474 and Treatment F_1,29_= 4.517, P = 0.0422, post-hoc: all P > 0.05 in Sham vs. ^56^Fe, ω_p_^2^= 0.04; interaction: BlockXTreatment F_2,58_= 0.8020, P = 0.4533) as well as the incorrect selection in the Large (**Fig. 3U, Table S1**, two-way RM ANOVA, main effect: Block F_2,58_ = 1.628, P = 0.2051 and Treatment F_1,29_ = 1.640, P = 0.2105; interaction: BlockXTreatment F_2,58_= 0.4078, P = 0.6670) and Small separation LDR Test blocks (**Fig. 3V Table S1**, two-way RM ANOVA, main effect: Block F_2,58_= 0.3911, P = 0.6781 and Treatment F_1,29_= 0.4058, P = 0.5291; interaction: BlockXTreatment F_2,58_= 0.6914, P = 0.5049). Together these data suggest ^56^Fe IRR mice have improved accuracy (yet increased impulsivity) in Large separation trials late in LDR Test, but no change in attention or motivation vs. Sham mice. ^56^Fe IRR mice also have increased impulsivity in Small separation trials late in LDR Test, but no change in performance. These results suggest ^56^Fe IRR mice are better than Sham IRR mice in key aspects of discrimination learning, despite showing impulsivity.

### 3.3. Whole-body ^56^Fe IRR does not change acquisition or extinction learning of a simple stimulus-response task

To determine whether the observed IRR-induced increase in cognitive performance was limited to hippocampal-dependent tasks, we next tested for PFC-dependent executive function. The same cohort of Sham and ^56^Fe IRR female mice underwent simple stimulus-response learning (acquisition; **Fig. 1; 4A**) followed by extinction of acquired learning (**Fig. 4F**). Stimulus-response learning was similar between the groups (**Fig. 4A**) as >75% of mice in both Sham and ^56^Fe IRR groups reached criteria by 60 days (**Fig. 4B; Table S1**) and both groups completed the task in a similar number of days (**Fig. 4C; Table S1**, t-test, t_(30)_ = 0.1179, P = 0.9069). In addition, when looking at general performance over the course of acquisition, Sham and ^56^Fe-IRR mice gave a similar number of correct responses to a stimulus (**Fig. 4E; Table S1**, two-way RM ANOVA, main effects: Session F_1,28_= 4.063, P = 0.0535 and Treatment F_1,28_= 2.071, P = 0.1612; interaction: SessionXTreatment F_1,28_= 2.073, P = 0.1610) in comparable times (**Fig. 4D; Table S1**, two-way RM ANOVA, main effects: Session F_1,28_= 50.16, P < 0.0001 and Treatment F_1,28_ = 0.4781, P = 0.4950; interaction of SessionXTreatment F_1,28_= 0.1246, P = 0.7267), again suggesting similar simple stimuli-response learning between groups.

In extinction testing **(Fig. 4F)**, Sham and ^56^Fe IRR mice took a similar number of days to reach criteria, indicating no difference in the rate of extinction learning (**Fig. 4H; Table S1**, t-test, t_(24)_ = 1.057, P = 0.3012). Sham and ^56^Fe IRR mice also had similar individual session length (**Fig. 4I; Table S1**, two-way RM ANOVA, main effect: Session F_2,50_= 87.44, P < 0.0001, ω_p_^2^=0.64 and Treatment F_1,25_= 1.077, P = 0.3092; interaction: SessionXTreatment F_2,50_ = 0.1365, P = 0.8727) and reached a stable omission criteria (>24 out of 30 response omissions) over the course of testing (**Fig. 4J; Table S1**, Mixed-effects analysis, main effects: Day of test F_23,559_ = 30.96, P < 0.0001 and Treatment F_1,25_= 0.05308, P = 0.8197; interaction: Day of testXTreatment F_23,559_= 1.101, P = 0.3389). To assess whether these same Touchscreen-experienced mice differed in measures of potential impulsivity or general engagement with the screen, we analyzed blank touches, blank touch latency, and ITI touches (**Fig. 4I-4M**). Sham and ^56^Fe IRR mice had a similar number of blank touches (**Fig.4K; Table S1**, two-way RM ANOVA, main effects: Session F_2,50_= 0.5277, P = 0.5932 and Treatment F_1,25_= 2.831, P = 0.1049; interaction: SessionXTreatment F_2,50_= 0.02480, P = 0.9755), speed to make a blank touch (**Fig. 4L; Table S1**, two-way RM ANOVA, main effect: Session F_2,50_= 9.867, P = 0.0002, ω_p_^2^ =0.17 and Treatment F_1,25_= 0.3933, P = 0.5363; interaction: SessionXTreatment F_2,50_= 1.324, P = 0.2752), and ITI touch’s number (**Fig. 4M; Table S1**, two-way RM ANOVA, main effects: Session F_2,50_ = 7.996, P = 0.0010, ω_p_^2^=0.11 and Treatment F_1,25_= 0.06728, P = 0.7975; interaction: SessionXTreatment F_2,50_= 0.5152, P = 0.6005). Together these results suggest no effect of ^56^Fe IRR on task-specific impulsivity. However, in the last extinction session, ^56^Fe IRR mice took ∼1.5 sec longer to give a correct response vs. Sham mice (**Fig. 4N; Table S1**, two-way RM ANOVA, main effects: Session F_2,50_= 3.835, P = 0.0282, ω_p_^2^=0.08 and Treatment F_1,25_= 1.512, P = 0.2302; interaction of SessionXTreatment F_2,50_ = 4.173, P = 0.0211, post-hoc: *a’* P = 0.0088 in Sham vs. ^56^Fe, ω_p_^2^ =0.08). This small but significant increase in latency for 56Fe IRR mice to give a correct response did not influence extinction performance. Therefore, taken together, these data suggest ^56^Fe IRR does not influence extinction performance.

### 3.4. Female mice given whole-body ^56^Fe IRR took twice as long as Sham IRR mice to reach stimulus-response habit learning criteria

It has been suggested that systems of “declarative” and “habit” memory relying upon the medial temporal lobe (e.g. hippocampus) and basal ganglia (e.g. caudate-putamen), respectively, may compete with one another during behavioral tasks^34^. They may be identifiably separable or function simultaneously, “overriding” one another during various learning tasks. To assess whether the observed IRR-induced increase in hippocampal-dependent discrimination learning occurs at the expense of striatal memory circuit functional integrity, a parallel group of mice was used to assess the influence of ^56^Fe IRR on visuomotor conditional learning (VMCL; **Fig. 1, 5**). VMCL reflects stimulus-response habit or “rule-based” learning and relies on intact circuits of the striatum and posterior cingulate cortex^31^.

Similar to what was seen with the parallel cohort of mice (**Fig. 2C**), during general touchscreen training this cohort of Sham and ^56^Fe IRR mice completed each training phase in a similar number of days (**Fig. 5A**, Mixed-effects analysis, main effects: Training phase F_4,71_= 116.1, P < 0.0001 and Treatment F_1,18_ = 0.04624, P = 0.8322; interaction: Training phaseXTreatment F_4,71_ = 1.573, P = 0.1908). Thus, in two parallel cohorts - one assessed 3-mon post-IRR and the other 1-mon post-IRR - there was no overt effect of ^56^Fe IRR on operant learning. Unfortunately in-depth accuracy analysis of the last stage (Punish Incorrect) was not possible due to computer file inaccessibility. In VMCL Train - an intermediate training phase prior to VMCL Test - Sham and ^56^Fe IRR mice also did not differ in completion days (26.27 vs. 25 days, respectively, **Fig. 5B,D; Table S1**). However, in the VMCL Test (**Fig. 5C**), ^56^Fe IRR mice took nearly twice as many days vs. Sham to reach criteria (**Fig. 5D**; **Table S1**, Mixed-effects analysis, main effects: Experiment phase F_1,18_ = 8.145, P = 0.0105 and Treatment F_1,18_= 6.334, P = 0.0215; interaction: Experiment phaseXTreatment F_1,18_= 7.329, P = 0.0144, post-hoc: *a’* P=0.0014 in Sham vs.^56^Fe in test). These data suggest a ^56^Fe IRR-induced impairment in the rate of striatal-mediated learning.

To assess whether a slower rate of VMCL learning in ^56^Fe IRR mice could be explained by behavioral deficits evident in earlier training stages, we analyzed VMCL Train performance in-depth (**Fig. 5B,E-I**). In VMCL Train, a similar proportion of Sham and ^56^Fe IRR mice reached criteria over time (50% subjects reached criteria at 31 days in both Sham and ^56^Fe IRR mice; **Fig. 5E; Table S1**, Log-rank [Mantel-Cox], P = 0.6512). Sham and ^56^Fe IRR mice also had similar length of training sessions (**Fig. 5F; Table S1**, two-way RM ANOVA, main effects: Session F_1,18_= 47.84, P < 0.0001, ω_p_^2^=0.55 and Treatment F_1,18_ = 3.606, P = 0.0737; interaction: SessionXTreatment F_1,18_ = 3.606, P = 0.0737), number of training trials (**Fig. 5G; Table S1**, two-way RM ANOVA, main effects: Session F_1,18_ = 107.9, P < 0.0001, ω_p_^2^=0.55 and Treatment F_1,18_= 1.034, P = 0.3227; interaction: SessionXTreatment F_1,18_ = 1.629, P = 0.2181), response accuracy (**Fig. 5H; Table S1**, two-way RM ANOVA, main effects: Session F_1,18_= 20.66, P = 0.0003, ω_p_^2^=-0.01 and Treatment F_1,18_ = 1.543, P = 0.2302; interaction: SessionXTreatment F_1,18_ = 0.4270, P = 0.5217), and number of correction trials following an incorrect response (**Fig. 5I; Table S1**, two-way RM ANOVA, main effects: Session F_1,18_= 9.162, P = 0.0072, ω_p_^2^ =0.19 and Treatment F_1,18_ = 1.928, P = 0.1819; interaction: SessionXTreatment F_1,18_= 1.611, P = 0.2205). Taken together, these results suggest no gross impact of ^56^Fe IRR on the ability to complete VMCL Train.

In VMCL Test (**Fig. 5C**), a similar distribution of the proportion of Sham and ^56^Fe IRR mice reach criteria over the entire VMCL Test period (**Fig. 5J; Table S1**, Log-rank [Mantel-Cox], P = 0.6501). Sham and ^56^Fe IRR mice had similar VMCL Test performance as indicated by similar session length (**Fig. 5K; Table S1**, Mixed-effects analysis, main effects: Session F_3,69_ = 1.318, P = 0.2755 and Treatment F_1,69_ = 2.212, P = 0.1415; interaction: SessionXTreatment F_3,69_= 0.7373, P = 0.5334), accuracy (**Fig. 5L; Table S1**, Mixed-effects analysis, main effects: Session F_3,51_= 1.878, P = 0.1450 and Treatment F_1,18_= 0.4497, P = 0.5110; and interaction of SessionXTreatment F_3,51_ = 0.9603, P = 0.4186), percentage of trials missed due to inactivity (**Fig. 5M; Table S1**, Mixed-effects analysis, main effects: Session F_3,51_ = 0.9020, P = 0.4467 and Treatment F_1,18_ = 0.2195, P = 0.6451; interaction: SessionXTreatment F_3,51_= 0.3523, P = 0.7876), and number of incorrect trials made during the initial choice stage (**Fig. 5N; Table S1**, main effects: Session F_3,51_= 1.016, P = 0.3931 and Treatment F_1,18_= 0.04586, P = 0.8328; interaction: SessionXTreatment F_3,51_ = 0.6279, P = 0.6003). Therefore, while ^56^Fe IRR mice took more days to complete VMCL Test at certain accuracy vs. Sham mice, this was not due to a difference in other VMCL training and testing performance measures.

### 3.5. Whole-body ^56^Fe IRR does not change measures relevant to anxiety, depression, repetitive behavior, and sociability

^56^Fe IRR-induced improvements in discrimination learning and the increased number of blank touches during LDR test could be explained by increased compulsivity or other stereotypic behaviors, or alterations in anxiety- or depair-like behaviors. To assess these possibilities, the same touchscreen-experienced Sham and ^56^Fe IRR mice were run on a variety of classic non-touchscreen behavior tests including elevated plus maze, marble burying,open field, social interaction, and forced swim test (**Fig. 1; 6A**).

To analyze and compare anxiety-like behavior between Sham and ^56^Fe IRR groups, mice were exposed to the elevated plus maze and open field, both well-validated anxiety tests in rodent models. In the elevated plus maze, Sham and ^56^Fe IRR mice spent a similar amount of time in both the open and closed arms (**Fig. 6B,C; Table S1**, unpaired t-test, t_(27)_ = 1.298, P = 0.2052 **[B]**, t_(27)_ = 1.229, P = 0.2298 **[C]**). In the open field, Sham and ^56^Fe IRR mice traveled a similar total distance and spent a similar amount of time in both predefined center and corner areas (**Fig. 6E-G; Table S1**, unpaired t-test, t_(27)_ = 0.6491, P = 0.5218 **[E]**, t_(27)_ = 1.327, P = 0.1958 **[F]**, t_(27)_ = 0.6805, P = 0.5020 **[G]**).

**Figure 6.**
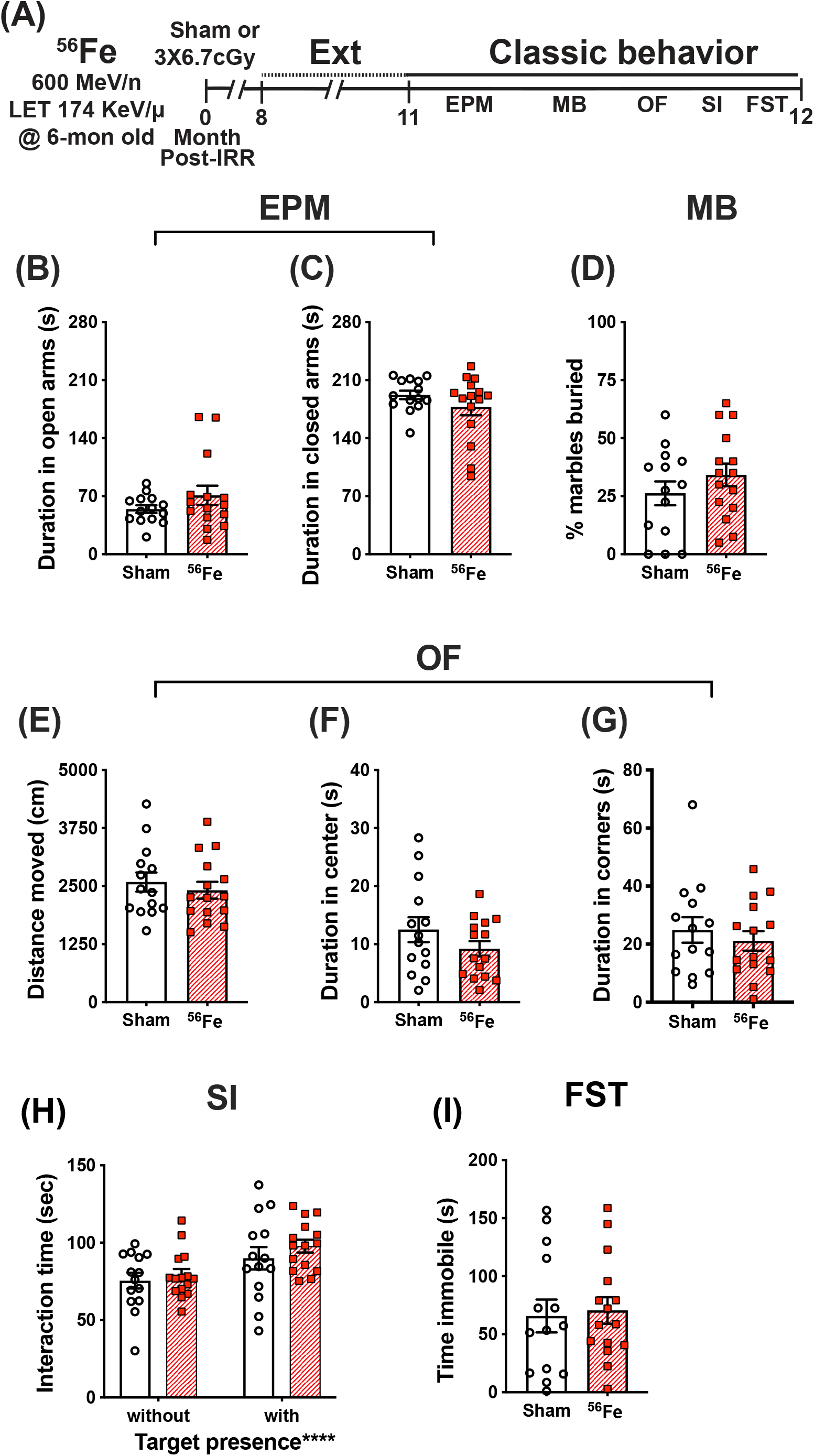
Timeline and results of classical behavior tests show anxiety-, compulsive-like behavior, and despair-like behavior and sociability are generally unaffected in female mice exposed to whole-body ^56^Fe IRR at 6-mon-old compared to Sham. **(A)** Six-month old C57BL/6J female mice that received whole-body exposure to ^56^Fe (0-Month Post-Irradiation [IRR]) and finished touchscreen LDR and Ext were then run on a variety of non-touchscreen classic behavioral tests. **(B**,**C)** Sham and ^56^Fe IRR mice showed no difference in anxiety-like behavior as shown by the time spent in either the open arms **(B)** or closed arms **(C)** of the elevated plus maze (EPM). **(D)** Sham and ^56^Fe IRR mice showed no difference in compulsive-like behavior as shown by percent of marbles buried in the marble burying (MB) task. **(E-G)** Sham and ^56^Fe IRR mice showed no difference in anxiety-like behavior as shown by the distance moved **(E)** and time spent in the center **(F)** or corner **(G)** of the open field (OF). **(H)** Sham and ^56^Fe IRR mice showed similar sociability when exposed to an age- and sex-matched conspecific in the social interaction (SI) test. Both groups spent more time in the interaction zone in the presence of a social target vs. in the absence of a social target. **(I)** Sham and ^56^Fe IRR mice had similar despair-like behavior as they spent a similar time immobile time in the forced swim test (FST). Error bars depict mean±SEM. Statistical analysis in **(H)**: Two-way RM ANOVA, main effect of target presence, ****p<0.0001. Bonferroni’s post-hoc analyses showed no difference between Sham and ^56^Fe IRR groups. Unpaired, two-tailed t-tests in **(B-G, I)**. cm, centimeters; EPM, elevated plus maze; Ext, extinction; FST, forced swim test; IRR, irradiation; LDR, location discrimination reversal; MB, marble burying; Mon; month; OF, open field; s, seconds; SI, social interaction.

To measure repetitive and compulsive-like behavior, the same female mice also were assessed in a marble burying test. Both Sham and ^56^Fe IRR mice buried a similar percentage of marbles during a 30-min session, which implies a lack of potentially pathological ^56^Fe IRR-induced stereotypic and compulsive behavior (**Fig. 6D; Table S1**, unpaired t-test, t_(27)_ = 1.119, P = 0.2729).

In certain transgenic rodent models of autism, mice show improved behavioral “pattern separation” alongside social deficits (Benevento et al., 2017). To assess whether ^56^Fe IRR-induced improvements in discrimination learning shown here were accompanied by social deficits, Sham and ^56^Fe IRR female mice were exposed to an age-, sex-matched conspecific in a social interaction test. Sham and ^56^Fe IRR mice spent a similar amount of time in a predefined interaction zone in the absence and presence of a conspecific target (in a plastic and wire mesh enclosure), and spent relatively more time in the interaction zone in the presence vs. absence of a conspecific (**Fig. 6H; Table S1**, two-way RM ANOVA, main effects: Target F_1,27_= 21.43, P < 0.0001, ω_p_^2^=0.15 and Treatment F_1.27_= 0.8003, P = 0.3789; interaction: TargetXTreatment F_1,27_= 0.3585, P = 0.5543). These data indicate no effect of ^56^Fe IRR on sociability.

We finally looked at whether ^56^Fe IRR-induced improvement in learning related to behavior shown under conditions that mimic despair (Lezak et al., 2017) by exposing the mice to the forced swim test. Both Sham and ^56^Fe IRR mice spent a similar amount of time immobile during the 6-min test (**Fig. 6I; Table S1**, unpaired t-test, t_(27)_ = 0.2643, P = 0.7936), suggesting no ^56^Fe IRR-induced despair-like phenotype.

## 4. Discussion

Here we provide a behavioral profile of female C57BL/6J mice that received a Mars mission-relevant dose of whole body ^56^Fe IRR at 6-mon of age. From 7-to 18-mon of age, ^56^Fe IRR mice and their Sham counterparts - which received every experimental manipulation but placement in front of the beam line - were examined via touchscreen and classical behavior tests to assess a range of cognitive abilities. We leveraged the power of the touchscreen platform to provide a holistic, multi-dimensional perspective on mouse behavior, performing in-depth analyses that have become the gold standard in the field (Horner et al., 2013; Oomen et al., 2013; Beraldo et al., 2019). Six differences emerged between Sham and ^56^Fe IRR mice. Relative to Sham mice, ^56^Fe IRR mice 1) took longer (27% longer) to complete the first session of the last stage of general training; 2) took ∼5s less (71% less time) to touch a blank window during the last stage of general training; 3) had a greater percent of correct trials (34% more) when distinguishing conditioned stimuli separated by a large (but not small) distance, specifically prior to their first reversal and late in location discrimination reversal testing; 4) touched a blank window more when distinguishing conditioned stimuli separated by a large or small distance (range of 43-63% more), specifically late in location discrimination reversal testing; 5) took ∼1s longer (34% longer) to touch the lit window (an incorrect response) in the last extinction session; and 6) took more than twice as many days (57% as many) to reach criteria in the visuomotor conditioned learning test. Sham and ^56^Fe IRR mice were similar in the other many touchscreen and classical behavior metrics collected. Below we discuss these findings in mice as they relate to cognitive domain function, indicate the strengths and limitations of our work, and speculate what these findings mean for NASA’s risk assessment for female astronauts in future deep space missions.

Our touchscreen data show that ^56^Fe IRR mice had ∼34% more correct trials relative to Sham mice, during Large separation trials prior to their first reversal and late in LDR testing. The effect size for this ^56^Fe IRR-induced increase in location discrimination is small (**Table S1**), and it is only seen in trials that use a Large, not Small, separation. In fact, both Sham and ^56^Fe IRR female mice perform just above chance (∼50% correct prior to first reversal, which is expected given the challenge of the task and the age of the mice at testing), and it is only in the last block that the ^56^Fe IRR mice perform better. This result suggests ^56^Fe IRR “improves” location discrimination reversal in female mice vs. Sham female mice. Given the well-described role of the hippocampus in LD and LDR (Clelland et al., 2009; McTighe et al., 2009; Swan et al., 2014), one interpretation of these data is that ^56^Fe IRR improves hippocampal function or perhaps integrity. There are three notable aspects of this interpretation. First, prior work with male mice reported that the same ^56^Fe IRR parameters used here improved discrimination learning in an LD test; LDR was not assessed in that study. In that study, ^56^Fe IRR male mice made a greater percentage of accurate responses and reached LD criteria in fewer days relative to Sham male mice. While there are additional distinctions between these studies, it is notable that both female and male mice exposed to ^56^Fe IRR show indices of improved LD or LDR - and thus perhaps improved hippocampal function - vs. Sham mice. Second, it is interesting to compare the interpretation of the present data (^56^Fe IRR female mice have improved LDR and hippocampal function) to prior literature on the impact of space radiation on hippocampal function. Many rodent studies suggest HZE particle exposure is detrimental to brain physiology and functional cognitive output, including hippocampal function (Blakely and Chang, 2007; Cucinotta et al., 2014; Kokhan et al., 2016; Nelson, 2016; Jandial et al., 2018; Cucinotta and Cacao, 2019; Kiffer et al., 2019; Limoli, 2020; Davis et al., 2021), although it is only recently that rodents have been irradiated at “astronaut age” or that female rodents have been more commonly studied. Indeed, the female rodent brain may be protected from radiation-induced immune and cognitive deficits (Villasana et al., 2011; Krukowski et al., 2018), or at least it appears there are sex-specific effects in the impact of ^56^Fe exposure on hippocampal-dependent function (Villasana et al., 2010, 2011). With these studies in mind, it is notable that touchscreen analysis of both female and male mice (of “astronaut age” at time of exposure) show ^56^Fe IRR improves LDR (present results) or LD (Whoolery et al., 2020), respectively, vs. Sham exposure without influencing other cognitive domains (exceptions in female mice are discussed below). A third perspective on these data with ^56^Fe IRR improving LDR in female mice is highlighted in recent work showing that the rodent brain has a compensatory, dynamic, time-dependent response to ^56^Fe IRR (Miry et al., 2021). More longitudinal studies are needed to clarify the time course of the LDR “improvement” reported here in ^56^Fe IRR female mice.

Another outcome of our touchscreen data is that ^56^Fe IRR female mice take more days to reach criteria relative to Sham mice in VMCL Test, suggesting impairment in stimulus-response habit learning. Given the reliance of rule-based learning on intact striatal circuits (Horner et al., 2013; Delotterie et al., 2015), our data suggest ^56^Fe IRR female mice have a deficit in striatal or basal ganglia function. Of note, while we did not observe behavioral changes that are indicative of gross striatal dysfunction (normal locomotor, marble burying, etc.), both high and low doses of HZE particles have been shown to produce maladaptive striatal plasticity and/or compromise the dopaminergic system of rodents (Joseph et al., 1992, 1993, 1994; Kiffer et al., 2019). Our finding that ^56^Fe IRR female mice have impaired stimulus-response habit learning is distinct from what is seen in male mice, as ^56^Fe IRR male mice perform similarly to Sham mice on VMCL (Whoolery et al., 2020). ^56^Fe IRR-induced deficits in stimulus-response habit learning in female mice is also opposite of our hypothesis, which was fueled by studies suggesting the female rodent brain may be spared from the negative impact of HZE particle exposure (Rabin et al., 2013; Krukowski et al., 2018). More specifically, when exposed to space radiation female mice - unlike male mice - do not show deficits in social interaction or novel social and object recognition memory, do not show anxiety-like phenotypes, and do not have the microglia activation and hippocampal synaptic losses seen in IRR male mice (Krukowski et al., 2018; Parihar et al., 2020). Thus, our data presented here add to the growing literature that whole-body exposure to HZE particles - such as ^56^Fe - affects cognition of mature male and female mice in a circuit-specific manner.

Taken together with our data presented here on the performance of female mice on LDR, it is notable that female mice perform *“*worse” on a striatal-dependent task, VMCL, but “better” on a hippocampal-dependent task, LDR. The present VMCL Test results are therefore interesting in regard to the theory of multiple memory systems (Nadel, 1992; Poldrack and Packard, 2003). Human and non-human memory studies suggest memory formation and consolidation are dependent on both hippocampal and non-hippocampal (i.e. basal ganglia or striatal) cooperative networks or memory systems. These systems encode for different memory types, with hippocampal circuits encoding relational memory for declarative past events and striatal circuits encode for acquisition of stimulus-response habit learning and some forms of Pavlovian conditioning (Poldrack and Packard, 2003). These networks also compete. In amnesic patients with partial temporal lobe damage, hippocampal-reliant recognition memory is decreased while striatal-mediated motor learning is spared (Tranel et al., 1994). Conversely, in patients with basal ganglia damage striatal memory function is decreased while hippocampal memory function is spared (Heindel et al., 1989). Here we report an ^56^Fe IRR-induced improvement in hippocampal-based discrimination learning yet deficits in striatal-dependent rule-based learning. We advocate for more specific evaluation of striatal-reliant behavioral patterns after HZE exposure using other touchscreen (i.e. autoshaping)(Horner et al., 2013) or other operant paradigms (rodent psychomotor vigilance test)(Davis et al., 2016), as such studies may clarify whether the improved discrimination learning shown here is accompanied by general basal ganglia-learning deficits. Additional study is also needed to determine if the results presented here in female mice - improved hippocampal-based LDR, decreased striatal-based VMCL - are a result of memory system competition.

In addition to the improved performance in LDR and decreased performance in VMCL, there are two other aspects of our data worth discussing. One, ^56^Fe IRR and Sham female mice in general performed similarly in Acquisition and Extinction, suggesting no difference in prefrontal cortical function. An exception is response latency; ^56^Fe IRR female mice take ∼1.5 s longer than Sham female mice to press the image. While this difference did not influence any other metric in Extinction, it is notable since the “correct” response in Extinction is to not touch the image. Here the ^56^Fe IRR female mice still press the image (which is an incorrect response) but take slightly longer to press it. Future studies will be needed to assess whether this longer latency means the ^56^Fe IRR mice are “in conflict” about making a response (but press it anyways). Two, there are indications that ^56^Fe IRR mice may be more impulsive. In the last and longest stage of general touchscreen training (PI), ^56^Fe IRR mice tested 3-mon post-IRR took 71% less time to touch blank windows in the last session vs. Sham mice. This is notable in that ^56^Fe IRR mice initially had 27% longer sessions early in PI. This faster blank touch in ^56^Fe IRR mice is not due to changes in attention, locomotor ability, or motivation since there are no differences between ^56^Fe IRR and Sham mice in the latency to touch the correct image or collect the reward. We interpret the shorter latency to touch a blank window in the last PI session 3-mon post-IRR as ^56^Fe IRR-induced impulsivity. It is unclear if the faster blank touch latency in ^56^Fe IRR mice late in PI is due to time post-IRR; we were unable to assess latency and other accuracy metrics in Sham and ^56^Fe IRR mice tested 1-mon post-IRR due to computer file issues. Another suggestion of impulsivity from our data is that ^56^Fe IRR mice touched blank windows more in both Large and Small separation trials near the end of LDR Test. As research suggests striatal circuits can also be involved in impulsive as well as habit behaviors (Fineberg et al., 2010; Lipton et al., 2019), it will be interesting for future space radiation studies to more specifically target assessment of impulsivity as it relates to striatal function and integrity.

The mechanism underlying ^56^Fe IRR-induced improvement in discrimination learning and decrement in rule-based learning is unknown, although the hippocampus and striatum, respectively, are linked to these functions (Clelland et al., 2009; McTighe et al., 2009; Horner et al., 2013; Oomen et al., 2013; Delotterie et al., 2015). Interestingly, a recent study reports ^56^Fe IRR induced hippocampal cellular, synaptic, and behavioral plasticity 2-mon post-IRR normalizes 6-mon post-IRR, and is actually enhanced 12-mon post-IRR (Miry et al., 2021). Thus, we hypothesize that the improved hippocampal-dependent discrimination learning and decreased striatal-based habit learning shown here are due to dynamic and compensatory processes post-IRR that are brain-region specific.

In conclusion, we have used a translationally-relevant rodent touchscreen battery to analyze the functional integrity of female mouse cognitive domains and associated brain circuits following exposure to the HZE particle ^56^Fe, a major component of space radiation that is a potential threat to the success of future crewed interplanetary missions. Our data in female mice: 1) suggest an IRR-induced competition between memory systems, as we see improved hippocampal-dependent memory and decreased striatal-dependent memory, 2) show that IRR induces sex-specific changes in cognition, 3) suggest the power that extensive multimodal behavioral analyses would have in helping standardize reporting of results from disparate behavioral experiments, and 4) underscore the importance of measuring multiple cognitive processes in preclinical space radiation risk studies, thereby preventing NASA’s risk assessments from being based on a single cognitive domain.

## 5. Data Availability Statement

Raw data will be placed in two archives: MouseBytes (https://www.mousebytes.ca/home) and NASA data portal (https://data.nas.nasa.gov/) and are also available on written request.

## 6. Ethics Statement

Human subjects: No

Animal subjects: Yes

Ethics statement: The study was approved by three Ethics committees (the Institutional Animal Care and Use Committees at the University of Texas Southwestern Medical Center [UTSW], Children’s Hospital of Philadelphia [CHOP], and Brookhaven National Laboratories [BNL]). Specifically, animal procedures and husbandry were in accordance with the National Institutes of Health Guide for the Care and Use of Laboratory Animals, and performed in IACUC-approved facilities at UT Southwestern Medical Center (UTSW, Dallas TX; AAALAC Accreditation #000673, PHS Animal Welfare Assurance D16-00296, Office of Laboratory Animal Welfare [OLAW] A3472-01), Children’s Hospital of Philadelphia (CHOP, Philadelphia, PA; AAALAC Accreditation #000427, PHS Animal Welfare Assurance D16-00280 [OLAW A3442-01]) and Brookhaven National Laboratories (BNL, Upton NY; AAALAC Accreditation #000048, PHS Animal Welfare Assurance D16-00067 [OLAW A3106-01]).

## 7. Author Contributions (based on <u>Project CRediT</u>) *initials listed via author order on study*

Conceptualization: IS, SY, CWW, AJE

Methodology: IS, SY, CWW, AJE

Software: YR

Validation: IS, SY, PLK, AJE

Formal Analysis: IS, SY, PLK

Investigation: IS, SY, CWW, FHT, RPR, MJD,

ADG Resources: SY, AJE

Data Curation: IS, SY, PLK

Writing, original draft: IS, SY, PLK, AJE

Writing, review and editing: IS, SY, PLK, FCK, AJE

Visualization: IS, SY, PLK, AJE

Supervision: SY, AMS, AJE

Projection Administration: SY, AJE

Funding Acquisition: SY, AMS, FCK, AJE

## 8. Acknowledgements

We thank many scientists for technical support and helpful conversations including Kyung Jin Ahn, Lyles Clark, Dr. Shibani Mukherjee, Dr. Guillermo Palchik, Dr. Ann Stowe, Vanessa Torres, Dr. Angela K. Walker, and Kielen Zuurbier. We thank staff members of the Brookhaven National Laboratory and the NASA Space Radiation Laboratory, including Adam Rusek (Physics team leader), MaryAnn Petry (animal support director), Peter Guida (organization and technical support director) as well as all of their team members who help make our experiments possible.

## 12. Supplementary materials

<u>Table S1, Detailed Statistical Results</u>

## 13. Current institutional addresses

IS:

RPR:

CWW:

YR:

MJD:

ADG:

AMS: Center for Advanced Translational Stroke Science, University of Kentucky, 3rd Floor BBSRB, 741 S. Limestone Street, Lexington, KY 40536

---

Supplementary Table 1 Statement on “Contribution to the Field”

