## Supplemental Table 1 for "Multi-domain touchscreen-based cognitive assessment of C57BL/6J female mice shows whole body exposure to ^56^Fe particle space radiation in maturity improves discrimination learning yet impairs stimulus-response habit learning"

**Table S1. Detailed statistical results.**

Bold text and ***, P<0.05. Italicized text, 0.05<P<0.1. *Magnitudes of Partial omega-squared (for RM two-way ANOVA): 0.01 small; 0.06 medium; 0.14 large. N/A not applicable

| Subject | Figure | n | Mean | | | | | | | | Statistics (variables) | Main Effect *P<0.5 **P<0.01 ***P<0.001 ****P<0.0001 | F Value | P value | Post hoc Test (Bonferroni) | Effect size (when RM two-way ANOVA p<0.05, partial omega-squared is calculated where ≤0.05 small, ≥0.06 medium, ≥0.14 large) |
|---|---|---|---|---|---|---|---|---|---|---|---|---|---|---|---|---|
| Weights | 2A | Sham: 16 | Post-IRR (wks) | 0 | 16 | 22 | 28 | 34 | 40 | 46 | Mixed-effects analysis | time**** | F (36, 1067) = 66.05 | **P<0.0001** | Sham vs. 56Fe: all P>0.05 | |
| | | | Sham | 25.4 | 29.19 | 28.53 | 30.26 | 31.18 | 31.09 | 36.52 | | treatment | F (1, 30) = 0.02450 | P=0.8767 | | |
| | | Frac 20 cGy: 16 | Frac 20 cGy | 24.92 | 29.91 | 28.83 | 29.94 | 30.92 | 30.76 | 38.3 | | interaction** | F (36, 1067) = 1.586 | **P=0.0161** | | |
| Touchscreen Locomotion | 2B | Sham: 16 | LM | | | | | | | | Unpaired t-test | NA | NA | P=0.6979 | NA | |
| | | | Sham | 199.3 ± 15.75 | | | | | | | | | | | | |
| | | Frac 20 cGy: 16 | Frac 20 cGy | 191.9 ± 10.29 | | | | | | | | | | | | |
| General Touchscreen Training w/six windows | 2C | Sham: 15 | General TS training | | | | | | | | Two-way RM ANOVA | interaction | F (5, 145) = 1.035 | P=0.3992 | NA | |
| | | | Training Stage | HAB1 | HAB2 | IT | MT | MI | PI | | | training stage**** | F (5, 145) = 59.27 | **P<0.0001** | | 0.62 |
| | | | Sham | 1 | 1 | 2.2 | 5.267 | 1 | 13.93 | | | treatment | F (1, 29) = 2.511 | P=0.1239 | | |
| | | Frac 20 cGy: 16 | Frac 20 cGy | 1 | 1 | 3.438 | 5.938 | 1.125 | 17.94 | | | subject | F (29, 145) = 1.048 | P=0.4105 | | |
| Punish incorrect Session length | 2D | | Session length (s) | | | | | | | | Two-way RM ANOVA | interaction | F (1, 29) = 2.345 | P=0.1365 | Sham vs. 56Fe: *a'* P = 0.0228 | |
| | | Sham: 15 | Session | First | Last | | | | | | | session | F (1, 29) = 0.5688 | P=0.4568 | | |
| | | | Sham | 1055 | 1217 | | | | | | | treatment* | F (1, 29) = 4.532 | **P=0.0419** | | 0.07 |
| | | Frac 20 cGy: 16 | Frac 20 cGy | 1340 | 1285 | | | | | | | subject | F (29, 29) = 1.355 | P=0.2092 | | |
| Punish incorrect Trial Number | 2E | | Trial # | | | | | | | | Two-way RM ANOVA | interaction | F (1, 29) = 3.742 | P=0.0629 | NA | |
| | | Sham: 15 | Session | First | Last | | | | | | | session | F (1, 29) = 3.742 | P=0.0629 | | |
| | | | Sham | 30 | 30 | | | | | | | treatment | F (1, 29) = 3.742 | P=0.0629 | | |
| | | Frac 20 cGy: 16 | Frac 20 cGy | 29 | 30 | | | | | | | subject | F (29, 29) = 1.000 | P=0.5000 | | |
| Punish incorrect Percent correct | 2F | | % correct | | | | | | | | Two-way RM ANOVA | interaction | F (1, 29) = 0.01728 | P=0.8963 | NA | |
| | | Sham: 15 | Session | First | Last | | | | | | | session**** | F (1, 29) = 149.5 | **P<0.0001** | | 0.69 |
| | | | Sham | 43.11 | 83.33 | | | | | | | treatment | F (1, 29) = 0.5868 | P=0.4499 | | |
| | | Frac 20 cGy: 16 | Frac 20 cGy | 40.78 | 80.15 | | | | | | | subject | F (29, 29) = 1.225 | P=0.2941 | | |
| Punish incorrect ITI Touch | 2G | | ITI Touch | | | | | | | | Two-way RM ANOVA | interaction | F (1, 29) = 1.138 | P=0.2949 | NA | |
| | | Sham: 15 | Session | First | Last | | | | | | | session**** | F (1, 29) = 49.58 | **P<0.0001** | | 0.44 |
| | | | Sham | 22.4 | 7.133 | | | | | | | treatment | F (1, 29) = 0.3334 | P=0.5681 | | |
| | | Frac 20 cGy: 16 | Frac 20 cGy | 21.5 | 10.25 | | | | | | | subject | F (29, 29) = 1.039 | P=0.4591 | | |
| Punish incorrect Correct touch latency | 2H | | Correct touch latency (s) | | | | | | | | Two-way RM ANOVA | interaction | F (1, 29) = 1.127 | P=0.2971 | NA | |
| | | Sham: 15 | Session | First | Last | | | | | | | session**** | F (1, 29) = 43.99 | **P<0.0001** | | 0.38 |
| | | | Sham | 10.5 | 5.71 | | | | | | | treatment | F (1, 29) = 0.2134 | P=0.6476 | | |
| | | Frac 20 cGy: 16 | Frac 20 cGy | 10.95 | 4.336 | | | | | | | subject | F (29, 29) = 1.349 | P=0.2127 | | |
| Punish incorrect Correct left touch latency | 2I | | Correct left touch latency (s) | | | | | | | | Two-way RM ANOVA | interaction | F (1, 29) = 0.001474 | P=0.9696 | NA | |
| | | Sham: 15 | Session | First | Last | | | | | | | session*** | F (1, 29) = 16.07 | **P=0.0004** | | 0.17 |
| | | | Sham | 10.75 | 6.398 | | | | | | | treatment | F (1, 29) = 2.078 | P=0.1601 | | |
| | | Frac 20 cGy: 16 | Frac 20 cGy | 8.846 | 4.572 | | | | | | | subject | F (29, 29) = 1.448 | P=0.1622 | | |
| Punish incorrect Correct right touch latency | 2J | | Correct right touch latency (s) | | | | | | | | Two-way RM ANOVA | interaction | F (1, 29) = 0.005288 | P=0.9425 | NA | |
| | | Sham: 15 | Session | First | Last | | | | | | | session*** | F (1, 29) = 15.34 | **P=0.0005** | | 0.17 |
| | | | Sham | 9.364 | 5.117 | | | | | | | treatment | F (1, 29) = 2.961 | P=0.0960 | | |
| | | Frac 20 cGy: 16 | Frac 20 cGy | 7.384 | 2.976 | | | | | | | subject | F (29, 29) = 1.174 | P=0.3345 | | |
| Punish incorrect Blank touch latency | 2K | | Blank touch latency (s) | | | | | | | | Two-way RM ANOVA | interaction | F (1, 29) = 3.234 | P=0.0825 | Sham vs. 56Fe: *a'*P = 0.0203 | |
| | | Sham: 15 | Session | First | Last | | | | | | | session** | F (1, 29) = 9.015 | **P=0.0055** | | 0.12 |
| | | | Sham | 11.65 | 10.13 | | | | | | | treatment* | F (1, 29) = 3.842 | **P=0.05** | | 0.05 |
| | | Frac 20 cGy: 16 | Frac 20 cGy | 11.4 | 5.338 | | | | | | | subject | F (29, 29) = 1.035 | P=0.4635 | | |
| Punish incorrect Reward latency | 2L | | Reward latency (s) | | | | | | | | Two-way RM ANOVA | interaction* | F (1, 29) = 5.175 | **P=0.0305** | Sham vs. 56Fe: all P>0.05 | 0.07 |
| | | Sham: 15 | Session | First | Last | | | | | | | session**** | F (1, 29) = 30.43 | **P<0.0001** | | 0.34 |
| | | | Sham | 2.281 | 1.668 | | | | | | | treatment | F (1, 29) = 0.1001 | P=0.7540 | | |
| | | Frac 20 cGy: 16 | Frac 20 cGy | 2.768 | 1.293 | | | | | | | subject | F (29, 29) = 0.8725 | P=0.6421 | | |
| LDR training % reaching criteria | 3B | Sham: 15 | LD train criteria completion curve | | | | | | | | Log-rank (Mantel-Cox) test | NA | NA | P=0.8480 | NA | |
| | | | Sham | Median: 5 | | | | | | | | | | | | |
| | | Frac 20 cGy: 16 | Frac 20 cGy | Median: 9 | | | | | | | | | | | | |
| LDR training Days to completion | 3C | Sham: 15 | Days to completion | | | | | | | | Unpaired t-test | NA | NA | P=0.9258 | NA | |
| | | | Sham | 6.133 ± 2.004 | | | | | | | | | | | | |
| | | Frac 20 cGy: 16 | Frac 20 cGy | 5.938 ± 0.7386 | | | | | | | | | | | | |
| LDR training Trial number | 3D | | Trial # | | | | | | | | Two-way RM ANOVA | interaction | F (1, 29) = 0.4779 | P=0.4949 | NA | |
| | | Sham: 15 | Session | First | Last | | | | | | | session | F (1, 29) = 1.549 | P=0.2233 | | |
| | | | Sham | 39.67 | 40.67 | | | | | | | treatment | F (1, 29) = 0.001294 | P=0.9716 | | |
| | | Frac 20 cGy: 16 | Frac 20 cGy | 38.31 | 41.81 | | | | | | | subject** | F (29, 29) = 2.566 | P=0.0067 | | |
| LDR training Percent correct to 1st reversal | 3E | Sham: 15 | % correct | | | | | | | | Two-way RM ANOVA | interaction | F (1, 29) = 8.180e-006 | P=0.9977 | NA | |
| | | | Session | First | Last | | | | | | | session | F (1, 29) = 0.9060 | P=0.3490 | | |
| | | | Sham | 65.91 | 70.49 | | | | | | | treatment | F (1, 29) = 3.713 | P=0.0638 | | |
| | | Frac 20 cGy: 16 | Frac 20 cGy | 55.59 | 60.19 | | | | | | | subject | F (29, 29) = 1.230 | P=0.2905 | | |
| LDR test Large separation session length | 3G | | Large separation session length (s) | | | | | | | | Two-way RM ANOVA | interaction | F (2, 58) = 0.002083 | P=0.9979 | NA | |
| | | Sham: 15 | Block | 1 | 4 | 6 | | | | | | block | F (2, 58) = 2.002 | P=0.1443 | | |
| | | | Sham | 1800 | 1800 | 1798 | | | | | | treatment | F (1, 29) = 0.002083 | P=0.9639 | | |
| | | Frac 20 cGy: 16 | Frac 20 cGy | 1800 | 1800 | 1799 | | | | | | subject | F (29, 58) = 1.000 | P=0.4857 | | |
| LDR test Small separation session length | 3H | | Small separation session length (s) | | | | | | | | Two-way RM ANOVA | interaction | F (2, 58) = 0.002083 | P=0.9979 | NA | |
| | | Sham: 15 | Block | 1 | 4 | 6 | | | | | | block | F (2, 58) = 2.002 | P=0.1443 | | |
| | | | Sham | 1800 | 1800 | 1800 | | | | | | treatment | F (1, 29) = 0.002083 | P=0.9639 | | |
| | | Frac 20 cGy: 16 | Frac 20 cGy | 1800 | 1800 | 1800 | | | | | | subject | F (29, 58) = 1.000 | P=0.4857 | | |
| LDR test Large separation trial # | 3I | | Large separation trial # | | | | | | | | Two-way RM ANOVA | interaction | F (2, 58) = 1.877 | P=0.1622 | NA | |
| | | Sham: 15 | Block | 1 | 4 | 6 | | | | | | block | F (2, 58) = 1.737 | P=0.1851 | | |
| | | | Sham | 43.2 | 36.67 | 40.8 | | | | | | treatment | F (1, 29) = 0.4793 | P=0.4943 | | |
| | | Frac 20 cGy: 16 | Frac 20 cGy | 39.69 | 41.75 | 49.75 | | | | | | subject**** | F (29, 58) = 3.552 | P<0.0001 | | |
| LDR test Small separation trial # | 3J | | Small separation trial # | | | | | | | | Two-way RM ANOVA | interaction | F (2, 58) = 0.07135 | P=0.9312 | NA | |
| | | Sham: 15 | Block | 1 | 4 | 6 | | | | | | block | F (2, 58) = 2.175 | P=0.1228 | | |
| | | | Sham | 33.87 | 35.87 | 39.33 | | | | | | treatment | F (1, 29) = 1.916 | P=0.1768 | | |
| | | Frac 20 cGy: 16 | Frac 20 cGy | 39.31 | 42.88 | 44.56 | | | | | | subject**** | F (29, 58) = 4.118 | P<0.0001 | | |
| LDR test Large separation percent correct to 1st reversal | 3K | | Large separation % correct to 1st reversal | | | | | | | | Two-way RM ANOVA | interaction* | F (2, 58) = 3.761 | **P=0.0291** | Sham vs. 56Fe: *a'* P = 0.0220 | 0.05 |
| | | Sham: 15 | Block | 1 | 4 | 6 | | | | | | block | F (2, 58) = 0.5836 | P=0.5611 | | |
| | | | Sham | 63.84 | 56.55 | 51.69 | | | | | | treatment | F (1, 29) = 1.230 | P=0.2765 | | |
| | | Frac 20 cGy: 16 | Frac 20 cGy | 58.97 | 57.54 | 69.31 | | | | | | subject | F (29, 58) = 1.413 | P=0.1308 | | |
| LDR test Small separation percent correct to 1st reversal | 3L | | Small separation % correct to 1st reversal | | | | | | | | Two-way RM ANOVA | interaction | F (2, 58) = 1.149 | P=0.3242 | NA | |
| | | Sham: 15 | Block | 1 | 4 | 6 | | | | | | block | F (2, 58) = 0.5919 | P=0.5566 | | |
| | | | Sham | 40.99 | 50.29 | 38.53 | | | | | | treatment | F (1, 29) = 2.469 | P=0.1269 | | |
| | | Frac 20 cGy: 16 | Frac 20 cGy | 48.89 | 48.97 | 51.79 | | | | | | subject | F (29, 58) = 1.124 | P=0.3452 | | |
| LDR test Large separation reversal # | 3M | | Large separation reversal # | | | | | | | | Two-way RM ANOVA | interaction | F (2, 58) = 1.034 | P=0.3619 | NA | |
| | | Sham: 15 | Block | 1 | 4 | 6 | | | | | | block | F (2, 58) = 2.559 | P=0.0861 | | |
| | | | Sham | 0.8667 | 1.267 | 0.9333 | | | | | | treatment | F (1, 29) = 0.1469 | P=0.7043 | | |
| | | Frac 20 cGy: 16 | Frac 20 cGy | 0.6875 | 1.25 | 1.375 | | | | | | subject | F (29, 58) = 1.369 | P=0.1537 | | |
| LDR test Small separation reversal # | 3N | | Small separation reversal # | | | | | | | | Two-way RM ANOVA | interaction | F (2, 58) = 0.03173 | P=0.9688 | NA | |
| | | Sham: 15 | Block | 1 | 4 | 6 | | | | | | block | F (2, 58) = 1.992 | P=0.1457 | | |
| | | | Sham | 0.2 | 0.6 | 0.3333 | | | | | | treatment | F (1, 29) = 1.777 | P=0.1928 | | |
| | | Frac 20 cGy: 16 | Frac 20 cGy | 0.5 | 0.8125 | 0.5625 | | | | | | subject | F (29, 58) = 1.516 | P=0.0888 | | |
| LDR test Large separation blank touch | 3O | | Large separation blank touch | | | | | | | | Two-way RM ANOVA | interaction | F (2, 58) = 0.9088 | P=0.4087 | Sham vs. 56Fe: a' P = 0.0232 | |
| | | Sham: 15 | Block | 1 | 4 | 6 | | | | | | block | F (2, 58) = 0.7784 | P=0.4639 | | |
| | | | Sham | 55.6 | 55.27 | 54.93 | | | | | | treatment* | F (1, 29) = 6.262 | **P=0.0182** | | 0.09 |
| | | Frac 20 cGy: 16 | Frac 20 cGy | 67.81 | 66.06 | 78.69 | | | | | | subject** | F (29, 58) = 2.094 | P=0.0084 | | |
| LDR test Small separation blank touch | 3P | | Small separation blank touch | | | | | | | | Two-way RM ANOVA | interaction | F (2, 58) = 0.1156 | P=0.8910 | Sham vs 56Fe: *a'* P = 0.0204 Block 4, *a'* P = 0.0292 Block 6 | |
| | | Sham: 15 | Block | 1 | 4 | 6 | | | | | | block | F (2, 58) = 2.185 | P=0.1216 | | |
| | | | Sham | 63.33 | 48.07 | 59 | | | | | | treatment** | F (1, 29) = 11.49 | **P=0.0020** | | 0.17 |
| | | Frac 20 cGy: 16 | Frac 20 cGy | 88.25 | 78.81 | 88.31 | | | | | | subject*** | F (29, 58) = 2.625 | P=0.0009 | | |
| LDR test Large separation reward latency | 3Q | | Large separation reward latency (s) | | | | | | | | Two-way RM ANOVA | interaction | F (2, 58) = 1.095 | P=0.3414 | NA | |
| | | Sham: 15 | Block | 1 | 4 | 6 | | | | | | block | F (2, 58) = 0.6710 | P=0.5151 | | |
| | | | Sham | 0.7607 | 0.97 | 0.878 | | | | | | treatment | F (1, 29) = 1.083 | P=0.3066 | | |
| | | Frac 20 cGy: 16 | Frac 20 cGy | 2.284 | 0.8806 | 0.9556 | | | | | | subject | F (29, 58) = 0.9787 | P=0.5122 | | |
| LDR test Small separation reward latency | 3R | | Small separation reward latency (s) | | | | | | | | Two-way RM ANOVA | interaction | F (2, 58) = 1.426 | P=0.2485 | NA | |
| | | Sham: 15 | Block | 1 | 4 | 6 | | | | | | block | F (2, 58) = 1.675 | P=0.1962 | | |
| | | | Sham | 0.6313 | 0.5907 | 0.5373 | | | | | | treatment | F (1, 29) = 1.971 | P=0.1709 | | |
| | | Frac 20 cGy: 16 | Frac 20 cGy | 2.311 | 0.6363 | 0.6406 | | | | | | subject | F (29, 58) = 0.9379 | P=0.5642 | | |
| LDR test Large separation | 3S | | Large separation correct image latency (s) | | | | | | | | Two-way RM ANOVA | interaction | F (2, 58) = 0.9595 | P=0.3891 | NA | |
| | | Sham: 15 | Block | 1 | 4 | 6 | | | | | | block | F (2, 58) = 1.298 | P=0.2808 | | |

**Table S1. Detailed statistical results.**

Bold text and ***, P<0.05. Italicized text, 0.05<P<0.1. *Magnitudes of Partial omega-squared (for RM two-way ANOVA): 0.01 small; 0.06 medium; 0.14 large. N/A not applicable

| Subject | Figure | n | Mean | | | | | | Statistics (variables) | Main Effect *P<0.5 **P<0.01 ***P<0.001 ****P<0.0001 | F Value | P value | Post hoc Test (Bonferroni) | Effect size (when RM two-way ANOVA p<0.05, partial omega-squared is calculated where ≤0.05 small, ≥0.06 medium, ≥0.14 large) |
|---|---|---|---|---|---|---|---|---|---|---|---|---|---|---|
| correct image latency | 3S | | Sham | 8.746 | 23.89 | 11.76 | | | Two-way RM ANOVA | treatment | F (1, 29) = 0.2292 | P=0.6357 | NA | |
| | | Frac 20 cGy: 16 | Frac 20 cGy | 7.699 | 12.25 | 17.06 | | | | subject | F (29, 58) = 1.040 | P=0.4380 | | |
| LDR test Small separation correct image latency | 3T | | Small separation correct image latency (s) | | | | | | Two-way RM ANOVA | interaction | F (2, 58) = 0.8020 | P=0.4533 | Sham vs. 56Fe: all P>0.05 | |
| | | Sham: 15 | Block | 1 | 4 | 6 | | | | block | F (2, 58) = 1.979 | P=0.1474 | | |
| | | | Sham | 26.37 | 17.31 | 12.21 | | | | treatment* | F (1, 29) = 4.517 | **P=0.0422** | | 0.04 |
| | | Frac 20 cGy: 16 | Frac 20 cGy | 12.11 | 9.445 | 9.084 | | | | subject | F (29, 58) = 1.208 | P=0.2655 | | |
| LDR test Large separation incorrect image latency | 3U | | Large separation incorrect image latency (s) | | | | | | Two-way RM ANOVA | interaction | F (2, 58) = 0.4078 | P=0.6670 | NA | |
| | | Sham: 15 | Block | 1 | 4 | 6 | | | | block | F (2, 58) = 1.628 | P=0.2051 | | |
| | | | Sham | 6.914 | 15.63 | 12.22 | | | | treatment | F (1, 29) = 1.640 | P=0.2105 | | |
| | | Frac 20 cGy: 16 | Frac 20 cGy | 5.971 | 8.786 | 9.316 | | | | subject | F (29, 58) = 1.050 | P=0.4259 | | |
| LDR test Small separation incorrect image latency | 3V | | Small separation incorrect image latency (s) | | | | | | Two-way RM ANOVA | interaction | F (2, 58) = 0.6914 | P=0.5049 | NA | |
| | | Sham: 15 | Block | 1 | 4 | 6 | | | | block | F (2, 58) = 0.3911 | P=0.6781 | | |
| | | | Sham | 12.58 | 13.54 | 12.93 | | | | treatment | F (1, 29) = 0.4058 | P=0.5291 | | |
| | | Frac 20 cGy: 16 | Frac 20 cGy | 14.63 | 8.693 | 10.21 | | | | subject | F (29, 58) = 1.391 | P=0.1417 | | |
| Acquisition Criteria completion curve | 4B | Sham: 16 | Acquisition criteria completion curve | | | | | | Log-rank (Mantel-Cox) test | NA | NA | P>0.9999 | NA | |
| | | | Sham | Median: 27 | | | | | | | | | | |
| | | Frac 20 cGy: 16 | Frac 20 cGy | Median: 37 | | | | | | | | | | |
| Acquisition Days to completion | 4C | Sham: 16 | Days to completion | | | | | | Unpaired t-test | NA | NA | P=0.9069 | NA | |
| | | | Sham | 19.56 ± 4.460 | | | | | | | | | | |
| | | Frac 20 cGy: 16 | Frac 20 cGy | 20.31 ± 4.534 | | | | | | | | | | |
| Acquisition Session length | 4D | | Session length (s) | | | | | | Two-way RM ANOVA | interaction | F (1, 28) = 0.1246 | P=0.7267 | NA | |
| | | Sham: 15 | Session | First | Last | | | | | session*** | F (1, 28) = 50.16 | P<0.0001 | | |
| | | | Sham | 1285 | 813.3 | | | | | treatment | F (1, 28) = 0.4781 | P=0.4950 | | |
| | | Frac 20 cGy: 15 | Frac 20 cGy | 1187 | 760.6 | | | | | subject** | F (28, 28) = 2.939 | P=0.0029 | | |
| Acquisition Correct response # | 4E | | Correct reponse # | | | | | | Two-way RM ANOVA | interaction | F (1, 28) = 2.073 | P=0.1610 | NA | |
| | | Sham: 15 | Session | First | Last | | | | | session | F (1, 28) = 4.063 | P=0.0535 | | |
| | | | Sham | 27.47 | 29.87 | | | | | treatment | F (1, 28) = 2.071 | P=0.1612 | | |
| | | Frac 20 cGy: 15 | Frac 20 cGy | 29.6 | 30 | | | | | subject | F (28, 28) = 1.286 | P=0.2553 | | |
| Ext Criteria completion curve | 4G | Sham: 15 | Extinction criteria completion curve | | | | | | Log-rank (Mantel-Cox) test | NA | NA | P=0.6830 | NA | |
| | | | Sham | Median: 18 | | | | | | | | | | |
| | | Frac 20 cGy: 15 | Frac 20 cGy | Median: 23 | | | | | | | | | | |
| Ext Days to completion | 4H | Sham: 15 | Days to completion | | | | | | Unpaired t-test | NA | NA | P=0.3012 | NA | |
| | | | Sham | 16.21 ± 1.867 | | | | | | | | | | |
| | | Frac 20 cGy: 15 | Frac 20 cGy | 13.58 ± 1.574 | | | | | | | | | | |
| Ext Session length | 4I | | Session length (s) | | | | | | Two-way RM ANOVA | interaction | F (2, 50) = 0.1365 | P=0.8727 | NA | |
| | | Sham: 13 | Session | 1 | 8 | Last | | | | session**** | F (2, 50) = 87.44 | **P<0.0001** | | 0.64 |
| | | | Sham | 506.2 | 567.5 | 580.9 | | | | treatment | F (1, 25) = 1.077 | P=0.3092 | | |
| | | Frac 20 cGy: 14 | Frac 20 cGy | 513 | 571.0 | 590.7 | | | | subject | F (25, 50) = 1.709 | P=0.0533 | | |
| Ext Omission # | 4J | | Omission # | | | | | | Mixed-effects analysis | | | | NA | |
| | | Sham: 13 | Session | 1 | 6 | 11 | 16 | 21 | | day of test**** | F (23, 559) = 30.96 | **P<0.0001** | | |
| | | | Sham | 12 | 20.43 | 22.64 | 24.71 | 24.03 | | treatment | F (1, 25) = 0.05308 | P=0.8197 | | |
| | | Frac 20 cGy: 14 | Frac 20 cGy | 12 | 20 | 22.38 | 24.15 | 24.7 | | interaction | F (23, 559) = 1.101 | P=0.3389 | | |
| Ext Blank touch | 4K | | Blank touch | | | | | | Two-way RM ANOVA | interaction | F (2, 50) = 0.02480 | P=0.9755 | NA | |
| | | Sham: 13 | Session | 1 | 8 | Last | | | | session | F (2, 50) = 0.5277 | P=0.5932 | | |
| | | | Sham | 14.85 | 17.08 | 16.08 | | | | treatment | F (1, 25) = 2.831 | P=0.1049 | | |
| | | Frac 20 cGy: 14 | Frac 20 cGy | 20.21 | 22.57 | 20.64 | | | | subject** | F (25, 50) = 2.741 | P=0.0012 | | |
| Ext Blank touch latency | 4L | | Blank touch latency (s) | | | | | | Two-way RM ANOVA | interaction | F (2, 50) = 1.324 | P=0.2752 | NA | |
| | | Sham: 13 | Session | 1 | 8 | Last | | | | session*** | F (2, 50) = 9.867 | **P=0.0002** | | 0.17 |
| | | | Sham | 2.882 | 3.818 | 4.569 | | | | treatment | F (1, 25) = 0.3933 | P=0.5363 | | |
| | | Frac 20 cGy: 14 | Frac 20 cGy | 3.145 | 3.706 | 3.926 | | | | subject | F (25, 50) = 1.314 | P=0.2024 | | |
| Ext ITI touch | 4M | | ITI Touch | | | | | | Two-way RM ANOVA | interaction | F (2, 50) = 0.5152 | P=0.6005 | NA | |
| | | Sham: 13 | Session | 1 | 8 | Last | | | | session*** | F (2, 50) = 7.996 | **P=0.0010** | | 0.11 |
| | | | Sham | 48.15 | 37.85 | 31.08 | | | | treatment | F (1, 25) = 0.06728 | P=0.7975 | | |
| | | Frac 20 cGy: 14 | Frac 20 cGy | 45.93 | 38.64 | 35.83 | | | | subject** | F (25, 50) = 2.302 | P=0.0060 | | |
| Ext Response latency | 4N | | Response latency (s) | | | | | | Two-way RM ANOVA | interaction* | F (2, 50) = 4.173 | **P=0.0211** | Sham vs. 56Fe: *a'* P = 0.0088 | 0.08 |
| | | Sham: 13 | Session | 1 | 8 | Last | | | | session* | F (2, 50) = 3.835 | **P=0.0282** | | 0.08 |
| | | | Sham | 4.54 | 5.150 | 4.753 | | | | treatment | F (1, 25) = 1.512 | P=0.2302 | | |
| | | Frac 20 cGy: 14 | Frac 20 cGy | 4.431 | 4.628 | 6.302 | | | | subject | F (25, 50) = 0.6457 | P=0.8813 | | |
| General Touchscreen Training w/three windows | 5A | | General TS training | | | | | | Mixed-effects analysis | | | | NA | |
| | | Sham: 11 | Training Stage | Hab 2 | IT | MT | MI | PI | | training phase**** | F (4, 71) = 116.1 | **P<0.0001** | | |
| | | Frac 20 cGy: 9 | Sham | 1.273 | 1.182 | 4.909 | 1.273 | 12 | | treatment | F (1, 18) = 0.04624 | P=0.8322 | | |
| | | | Frac 20 cGy | 1.222 | 1 | 4.222 | 2.873 | 10.89 | | interaction | F (4, 71) = 1.573 | P=0.1908 | | |
| VMCL train/test | 5D | | VMCL train/test | | | | | | Mixed-effects analysis | experiment phase* | F (1, 18) = 8.145 | **P=0.0105** | Sham vs. 56Fe: a' P = 0.0014 | |
| | | Sham: 11 | Experiment phase | Train | Test | | | | | | | | | |
| | | | Sham | 26.27 | 26.73 | | | | | treatment* | F (1, 18) = 6.334 | **P=0.0215** | | |
| | | Frac 20 cGy: 9 | Frac 20 cGy | 25 | 42.22 | | | | | interaction* | F (1, 18) = 7.329 | **P=0.0144** | | |
| VMCL train criteria completion curve | 5E | Sham: 11 | VMCL train criteria completion curve | | | | | | Log-rank (Mantel-Cox) test | NA | NA | P=0.6512 | NA | |
| | | | Sham | Median: 31 | | | | | | | | | | |
| | | Frac 20 cGy: 9 | Frac 20 cGy | Median: 31 | | | | | | | | | | |
| VMCL train Session length | 5F | | Session length (s) | | | | | | Two-way RM ANOVA | interaction | F (1, 18) = 3.606 | P=0.0737 | NA | |
| | | Sham: 11 | Session | First | Last | | | | | session**** | F (1, 18) = 47.84 | **P<0.0001** | | 0.55 |
| | | | Sham | 1800 | 1425 | | | | | treatment | F (1, 18) = 3.606 | P=0.0737 | | |
| | | Frac 20 cGy: 9 | Frac 20 cGy | 1800 | 1587 | | | | | subject | F (18, 18) = 1.000 | P=0.5000 | | |
| VMCL train Trial # | 5G | | Trial # | | | | | | Two-way RM ANOVA | interaction | F (1, 18) = 1.629 | P=0.2181 | NA | |
| | | Sham: 11 | Session | First | Last | | | | | session**** | F (1, 18) = 107.9 | **P<0.0001** | | 0.55 |
| | | | Sham | 12.91 | 25 | | | | | treatment | F (1, 18) = 1.034 | P=0.3227 | | |
| | | Frac 20 cGy: 9 | Frac 20 cGy | 15.22 | 24.67 | | | | | subject | F (18, 18) = 0.8817 | P=0.6039 | | |
| VMCL train Percent correct | 5H | | % correct | | | | | | Two-way RM ANOVA | interaction | F (1, 18) = 0.4270 | P=0.5217 | NA | |
| | | Sham: 11 | Session | First | Last | | | | | session*** | F (1, 18) = 20.66 | **P=0.0003** | | -0.01 |
| | | | Sham | 77.68 | 91.27 | | | | | treatment | F (1, 18) = 1.543 | P=0.2302 | | |
| | | Frac 20 cGy: 9 | Frac 20 cGy | 71.99 | 90.14 | | | | | subject | F (18, 18) = 0.6181 | P=0.8417 | | |
| VMCL train Correction trial # | 5I | | Correction trial # | | | | | | Two-way RM ANOVA | interaction | F (1, 18) = 1.611 | P=0.2205 | NA | |
| | | Sham: 11 | Session | First | Last | | | | | session** | F (1, 18) = 9.162 | **P=0.0072** | | 0.19 |
| | | | Sham | 3.909 | 2.545 | | | | | treatment | F (1, 18) = 1.928 | P=0.1819 | | |
| | | Frac 20 cGy: 9 | Frac 20 cGy | 5.889 | 2.556 | | | | | subject | F (18, 18) = 0.8527 | P=0.6305 | | |
| VMCL test Criteria completion curve | 5J | Sham: 11 | VMCL test criteria completion curve | | | | | | Log-rank (Mantel-Cox) test | NA | NA | P = 0.6501 | NA | |
| | | | Sham | Median: 36 | | | | | | | | | | |
| | | Frac 20 cGy: 9 | Frac 20 cGy | Median: 47 | | | | | | | | | | |
| VMCL test Session length | 5K | | Session length (s) | | | | | | Mixed-effects analysis | session | F (3, 69) = 1.318 | P=0.2755 | NA | |
| | | Sham: 11 | Session | 1 | 8 | 13 | 22 | | | | | | | |
| | | | Sham | 1800 | 1800 | 1754 | 1737 | | | treatment | F (1, 69) = 2.212 | P=0.1415 | | |
| | | Frac 20 cGy: 9 | Frac 20 cGy | 1800 | 1800 | 1800 | 1786 | | | interaction | F (3, 69) = 0.7373 | P=0.5334 | | |
| VMCL test Percent correct | 5L | | % correct | | | | | | Mixed-effects analysis | session | F (3, 51) = 1.878 | P=0.1450 | NA | |
| | | Sham: 11 | Session | 1 | 8 | 13 | 22 | | | | | | | |
| | | | Sham | 40.57 | 49.59 | 56.33 | 61.52 | | | treatment | F (1, 18) = 0.4497 | P=0.5110 | | |
| | | Frac 20 cGy: 9 | Frac 20 cGy | 50 | 36.7 | 50.05 | 54.53 | | | interaction | F (3, 51) = 0.9603 | P=0.4186 | | |
| VMCL test Percent missed | 5M | | % missed | | | | | | Mixed-effects analysis | session | F (3, 51) = 0.9020 | P=0.4467 | NA | |
| | | Sham: 11 | Session | 1 | 8 | 13 | 22 | | | | | | | |
| | | | Sham | 30.63 | 19.78 | 21.79 | 17.08 | | | treatment | F (1, 18) = 0.2195 | P=0.6451 | | |
| | | Frac 20 cGy: 9 | Frac 20 cGy | 26.36 | 27.05 | 26.95 | 19.42 | | | interaction | F (3, 51) = 0.3523 | P=0.7876 | | |
| VMCL test Incorrect trial # | 5N | | Incorrect trial # | | | | | | Mixed-effects analysis | session | F (3, 51) = 1.016 | P=0.3931 | NA | |
| | | Sham: 11 | Session | 1 | 8 | 13 | 22 | | | | | | | |
| | | | Sham | 4 | 3.182 | 4.182 | 4.657 | | | treatment | F (1, 18) = 0.04586 | P=0.8328 | | |
| | | Frac 20 cGy: 9 | Frac 20 cGy | 4 | 4.222 | 3.667 | 4.778 | | | interaction | F (3, 51) = 0.6279 | P=0.6003 | | |
| Elevated Plus Maze Open arm time | 6B | Sham: 14 | Open arm time (s) | | | | | | Unpaired t-test | NA | NA | P=0.2052 | NA | |
| | | | Sham | 54.48 ± 4.561 | | | | | | | | | | |
| | | Frac 20 cGy: 15 | Frac 20 cGy | 71.07 ± 11.58 | | | | | | | | | | |
| Elevated Plus Maze Closed arm time | 6C | Sham: 14 | Closed arm time (s) | | | | | | Unpaired t-test | NA | NA | P=0.2298 | NA | |
| | | | Sham | 192.1 ± 5.199 | | | | | | | | | | |
| | | Frac 20 cGy: 15 | Frac 20 cGy | 177.8 ± 10.20 | | | | | | | | | | |
| Marble Burying Percent marbles buried | 6D | Sham: 14 | % marbles buried | | | | | | Unpaired t-test | NA | NA | P=0.2729 | NA | |
| | | | Sham | 26.25 ± 5.161 | | | | | | | | | | |
| | | Frac 20 cGy: 15 | Frac 20 cGy | 34.17 ± 4.847 | | | | | | | | | | |
| Open Field Distance moved | 6E | Sham: 14 | Distance moved (cm) | | | | | | Unpaired t-test | NA | NA | P=0.5218 | NA | |
| | | | Sham | 2589 ± 205.3 | | | | | | | | | | |
| | | Frac 20 cGy: 15 | Frac 20 cGy | 2411 ± 180.9 | | | | | | | | | | |
| Open Field Center area time | 6F | Sham: 14 | Center area time (s) | | | | | | Unpaired t-test | NA | NA | P=0.1958 | NA | |
| | | | Sham | 12.50 ± 2.163 | | | | | | | | | | |
| | | Frac 20 cGy: 15 | Frac 20 cGy | 9.203 ± 1.303 | | | | | | | | | | |
| Open Field | | Sham: 14 | Corner area time (s) | | | | | | | | | | | |

**Table S1. Detailed statistical results.**

Bold text and ***, P<0.05. Italicized text, 0.05<P<0.1. *Magnitudes of Partial omega-squared (for RM two-way ANOVA): 0.01 small; 0.06 medium; 0.14 large. N/A not applicable

| Subject | Figure | n | Mean | | | Statistics (variables) | Main Effect<br>*P<0.5<br>**P<0.01<br>***P<0.001<br>****P<0.0001 | F Value | P value | Post hoc Test (Bonferroni) | Effect size (when RM two-way ANOVA p<0.05, partial omega-squared is calculated where ≤0.05 small, ≥0.06 medium, ≥0.14 large) |
|---|---|---|---|---|---|---|---|---|---|---|---|
| Open Field Corner area time | 6G | Sham: 14 | Sham | 24.90 ± 4.424 | | Unpaired t-test | NA | NA | P=0.5020 | NA | |
| | | Frac 20 cGy: 15 | Frac 20 cGy | 21.14 ± 3.376 | | | | | | | |
| Social interaction Interaction time | 6H | | Interaction time (s) | | | Two-way RM ANOVA | interaction | F (1, 27) = 0.3585 | P=0.5543 | NA | 0.15 |
| | | Sham: 14 | Target presence | without | with | | target**** | F (1, 27) = 21.43 | **P<0.0001** | | |
| | | | Sham | 75.46 | 89.91 | | treatment | F (1, 27) = 0.8003 | P=0.3789 | | |
| | | Frac 20 cGy: 15 | Frac 20 cGy | 79.05 | 97.79 | | subject** | F (27, 27) = 3.200 | P=0.0018 | | |
| Forced Swim Test Immobile time | 6I | Sham: 14 | Immobile time (s) | | | Unpaired t-test | NA | NA | P=0.7936 | NA | |
| | | | Sham | 65.72 ± 14.20 | | | | | | | |
| | | Frac 20 cGy: 15 | Frac 20 cGy | 70.51 ± 11.47 | | | | | | | |
